# Rhythmic modulation of subthalamo-pallidal interactions depends on synaptic rewiring through inhibitory plasticity

**DOI:** 10.1101/2024.07.01.601477

**Authors:** Mojtaba Madadi Asl, Caroline A. Lea-Carnall

## Abstract

Rhythmic stimulation offers a paradigm to modulate brain oscillations and, therefore, influence brain function. A growing body of evidence indicates that reciprocal interactions between the neurons of the subthalamic nucleus (STN) and globus pallidus externus (GPe) play a central role in the emergence of abnormal synchronous beta (15-30 Hz) oscillations in Parkinson’s disease (PD). The proliferation of inhibitory GPe-to-STN synapses following dopamine loss exacerbates this pathological activity. Rhythmic modulation of the STN and/or GPe, for example, by deep brain stimulation (DBS), can restore physiological patterns of activity and connectivity. Here, we tested whether dual targeting of STN-GPe by rhythmic stimulation can modulate pathologically strong GPe-to-STN synapses through inhibitory spike-timing-dependent plasticity (iSTDP). More specifically, we examined how time-shifted paired stimuli delivered to the STN and GPe can lead to inter-population synaptic rewiring. To that end, we first theoretically analysed the optimal range of stimulation time shift and frequency for effective synaptic rewiring. Then, as a minimal model for generating subthalamo-pallidal oscillations in healthy and PD conditions, we considered a biologically inspired STN-GPe loop comprised of conductance-based spiking neurons. Consistent with the theoretical predictions, rhythmic stimulation with appropriate time shift and frequency modified GPe-to-STN interactions through iSTDP, i.e., by long-lasting rewiring of pathologically strong synaptic connectivity. This ultimately caused desynchronising after-effects within each population such that excessively synchronous beta activity in the PD state was suppressed, resulting in a decoupling of the STN-GPe network and restoration of healthy dynamics in the model. Decoupling effects of the dual STN-GPe stimulation can be realised by time-shifted continuous and intermittent stimuli, as well as monopolar and bipolar simulation waveforms. Our findings demonstrate the critical role of neuroplasticity in shaping long-lasting stimulation effects and may contribute to the optimisation of a variety of multi-site stimulation paradigms aimed at reshaping dysfunctional brain networks by targeting plasticity.

## 1 Introduction

Abnormal oscillatory activity is a hallmark of several brain disorders such as Parkinson’s disease (PD), epilepsy, tinnitus, and schizophrenia [1–5]. In PD, for instance, interactions between reciprocally connected neurons of the subthalamic nucleus (STN) and globus pallidus externus (GPe) form a pacemaker in the basal ganglia [6], playing a key role in the emergence and maintenance of overly synchronised beta band (15-30 Hz) oscillations [7], as shown both experimentally [8–10] and computationally [11–14]. Excessive beta synchrony is related to the severity of motor symptoms in PD [15, 16], and is reduced by dopamine-based medication [17, 18] and deep brain stimulation (DBS) [19–21].

High-frequency (> 100 Hz) DBS is the standard therapy for medically refractory movement disorders such as PD [20, 22]. However, the underlying mechanism is not sufficiently understood [23]. Remarkably, parkinsonian symptoms such as tremor, bradykinesia, and rigidity return within minutes to a few hours when DBS is turned off [24], demanding chronic stimulation which may induce side-effects [25]. As such, treatments that provide long-lasting effects that persist after discontinuation of stimulation are a current clinical need. Our group has shown using *in silico* simulations that aberrant neuronal synchrony can be effectively counteracted by desynchronising stimulation [26, 27], and that long-lasting desynchronisation may depend on the reshaping of pathological synaptic connectivity [28–30].

Abnormal rhythmogenesis in the STN-GPe circuit is associated with maladaptive and/or compensatory changes in neuronal activity and synaptic connectivity following dopamine loss [31,32] (see Ref. [33] for a review). In particular, pallido-subthalamic synaptic transmission is augmented in 6-hydroxydopamine (6-OHDA)-lesioned rodents [34,35], accompanied by an increase in the number of GPe→ STN synaptic connections and their strengths as compared to healthy animals [34]. This augmented synaptic transmission was attributed to plasticity mechanisms triggered by the imbalance between glutamatergic cortico-subthalamic inputs and gammaaminobutyric acid (GABA)ergic pallido-subthalamic inputs to the STN [35], which may further intensify abnormal, correlated STN-GPe activity [34, 35].

Computational models can shed light on the mechanisms underlying abnormal STN-GPe interactions. The strength of inhibitory input from the striato-pallidal pathway to the STN turns out to be a key parameter controlling oscillations in the STN-GPe loop, as suggested by network models constrained by experimental data [36–38]. As shown recently, for example, a simple model of the STN-GPe network with inhibitory spike-timing-dependent plasticity (iSTDP) [39–41] can explain pathological strengthening of the GPe→ STN synapses as well as associated excessive STN-GPe beta synchronisation in the PD state [42]. Such plastic changes of synaptic connectivity are likely to serve as a potential mechasim controlling transitions between qualitatively different multistable dynamical states [43–46], e.g., transitions between healthy and PD dynamics as demonstrated experimentally [34,35] as well as computationally [37,42]. In fact, accumulating experimental evidence suggests that the STN-GPe interactions are mediated by synaptic plasticity [34, 35, 47, 48].

This is relevant not only from a physiological standpoint, but also for therapeutic purposes such as the development of model-based stimulation paradigms to counteract abnormal synchronisation through synaptic rewiring [29, 30]. These techniques would be designed to drive a reduction in pathologically strong synaptic weights, enabling long-lasting effects that outlast the cessation of stimulation [27]. In principle, synaptic rewiring can be achieved by applying stimuli to induce a time shift between neuronal spikes and, thereby, rewiring neuronal networks through plasticity by up- and down-regulation of the synaptic weights [30]. This was motivated by theory-based coordinated reset of neuronal subpopulations aimed at desynchronisation of networks by multi-site stimulation (i.e., targeting two or more populations) [26, 27], as validated in pre-clinical as well as clinical proof-of-concept studies [49–52]. By considering synaptic plasticity in the STN-GPe network models, coordinated reset stimulation was able to reduce neuronal spike coincidences and, hence, down-regulate abnormally strong synaptic weights when STN → STN [53] or GPe → GPe [37] connections were plastic.

Network topology is known to impact STN-GPe synchronisation [54]. Yet, to what extent changes in synaptic connectivity influence parkinsonian STN-GPe interactions has remained overlooked. As shown both experimentally and computationally, selective modulation of interpopulation connections relies on synaptic rewiring by paired simulation-induced Hebbian plasticity between the two populations [55, 56], implying that both pre- and postsynaptic populations must be targeted and engaged by stimulation. Therefore, we hypothesised that modulation of GPe→ STN synapses by dual STN-GPe stimulation may engage plasticity and favourably reshape STN-GPe interactions, promoting long-lasting stimulation effects. We tested our idea in a biologically plausible model of the STN-GPe loop where the strength of the inter-population GPe → STN synapses were modified by iSTDP.

More specifically, we established two qualitatively different model states by adjusting the external currents applied to the STN and GPe in our model. Desynchronised states with weak connectivity (modelling healthy states) as opposed to synchronised network states with strong connectivity (modelling PD states) were stable in the absence of stimulation. We then stimulated the STN and GPe with a specific frequency guided by theoretical predictions. The stimulation signal was delivered simultaneously to all cells within each target population, but with a theoretically predicted time shift (delay) [30] between the two populations to harness inhibitory plasticity specifically for the down-regulation of the GPe→ STN synaptic strengths. Our results show that appropriate stimulation time shift and frequency lead to effective synaptic rewiring between the two populations. Down-regulation of the inter-population synapses ultimately decoupled the STN and GPe, resulting in an uncorrelated STN-GPe network activity and suppression of excessive beta oscillations.

Our study demonstrates that dual STN-GPe targeting by time-shifted stimulation [30] can induce potent anti-kindling effects [27] by unlearning pathologically strong synaptic interactions, thus shifting the parkinsonian STN-GPe network characterised by strong synchronisation and strong connectivity to a more physiologically favoured, loosely connected, desynchronised state. Decoupling effects can be realised by time-shifted continuous and intermittent stimuli, as well as monopolar and bipolar simulation waveforms, suggesting that the dual STN-GPe stimulation paradigm might work in a more generic manner. The results of our modelling study might have strong implications in various multi-channel brain stimulation techniques aimed at reshaping dysfunctional diseased brain networks by targeting plasticity.

## 2 Methods

### 2.1 Network model

As a minimal model for generating subthalamo-pallidal oscillations, we considered a reciprocally connected STN-GPe network, schematically shown in Fig. 1A and B. In the model, the GPe provides inhibitory input to the STN (red solid line with circle) and, in turn, the STN excites the GPe (green dashed line with arrow). For simplicity, the striatal inhibition to the GPe (violet dashed line with circle) and cortical excitation to the STN (violet dashed line with arrow) were simulated as independent external currents (*I*_ext_).

**Figure 1:**
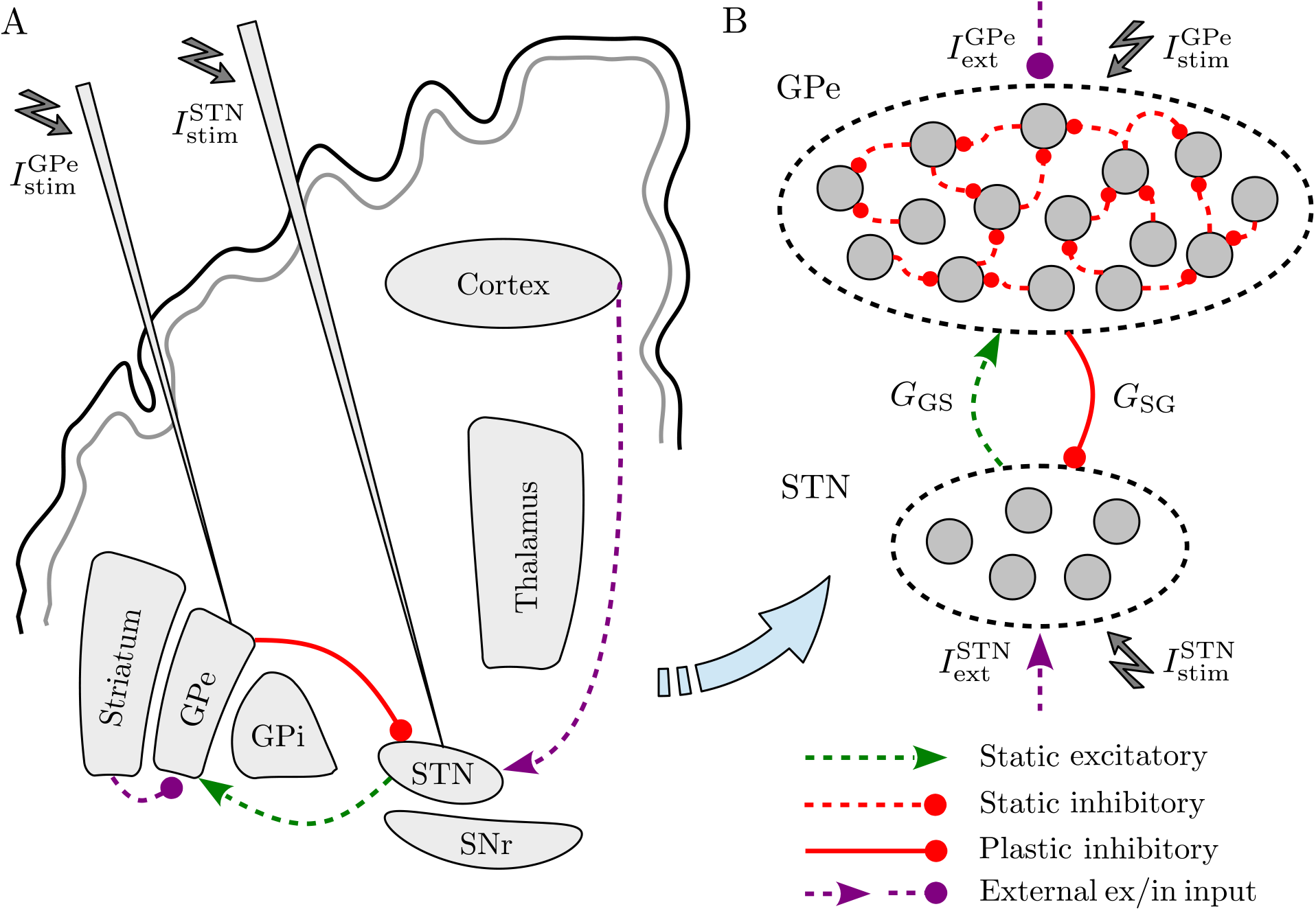
Schematics of the STN-GPe network model. (**A**) Overview of the model projections and the concept of the dual STN-GPe stimulation. (**B**) The reciprocally connected STN-GPe network with plastic GPe → STN synapses (red solid line with circle). Inter-population STN → GPe (GPe → STN) interactions are characterised by the mean coupling strength *G*_GS_(*G*_SG_). The striatal and cortical projections (violet dashed lines) are simulated as independent external currents (*I*_ext_) applied to the STN and GPe, and *I*_stim_ represents the corresponding stimulation current.

Physiological estimates of the number of basal ganglia neurons in rodents revealed that the GPe (∼46,000 cells) contains more than three times as many neurons as the STN (∼13,600 cells) [57]. Inspired by this experimental evidence, in our model the GPe contained 460 cells, whereas the STN was composed of 136 cells, such that the ratio of the cells was scaled by 0.01 (reduced to 1%) of their physiological estimates.

The STN-GPe network interactions were realised by random, sparsely connected GPe → GPe (connection probability 5%), GPe→ STN (connection probability 2%) and STN → GPe (connection probability 5%) pathways. Motivated by experimental evidence indicating a lack of intrinsic connectivity within the STN in rodents [58], the STN → STN connections were ignored in the model, i.e., the STN neurons were assumed to be independent elements driven by upstream structures. Connection probability and the range of synaptic strengths in the model are given in Table 1, which were inspired by experimental estimates in rodents [59–62] as well as relevant computational studies [11, 36, 63].

**Table 1:** Connection probability (*p*) [59–62] and the strength of synaptic connections (*g*) [11, 36, 62, 63] between the STN and GPe used in the model.

| Connection | $p$ | $g^*$ (mean) |
| --- | --- | --- |
| GPe → GPe | 0.05 | 0.25 |
| GPe → STN | 0.02 | 0.25 |
| STN → GPe | 0.05 | 0.25 |

\*Units in nS/ $\mu\text{m}^2$ .

## 2.2 Neuron model

The membrane potential dynamics of the STN/GPe neurons (*V*) in the loop was described by a single-compartment conductance-based model introduced by Terman and colleagues [11], as follows:

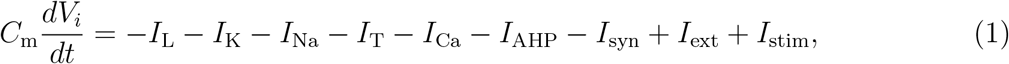

where *C*_m_ is the membrane capacitance, and *I*_ext_ represents the excitatory/inhibitory external current applied to the STN/GPe. In the model, the leak current (*I*_L_), potassium current (*I*_K_), sodium current (*I*_Na_) and high-threshold calcium current (*I*_Ca_) are described by the following Hodgkin-Huxley-type equations:

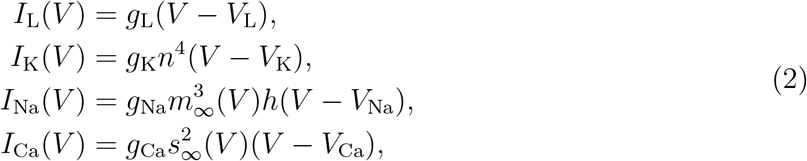

which are identical for the STN and GPe cells. The low-threshold T-type calcium current (*I*_T_) is defined differently for the STN and GPe neurons, as follows:

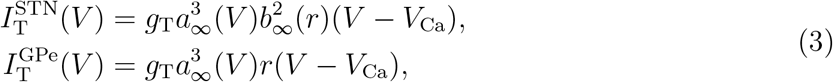

where *g*_X_ and *V*_X_ with X∈ {L, K, Na, Ca} represent the maximal conductance and reversal potential of each current, respectively. The first-order kinetics of slowly operating gating variables (i.e., *n, h, r*) can be described by the following differential equation:

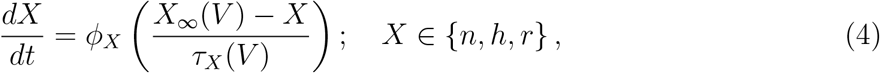

where ϕ_*X*_ is the scaling time constant of the variable *X*. The voltage-dependent time constant of the variable *X* can be written as follows:

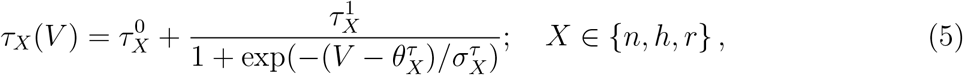

where 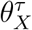represents the voltage at which the time constant is midway between its maximum and minimum values, and 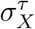 is the slope factor for the voltage dependence of the time constant. The steady-state voltage dependence of all gating variables is defined as follows:

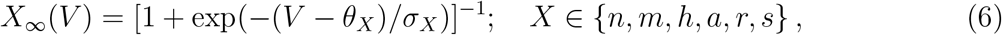

where*θ*_*X*_ is the half activation/inactivation voltage and *σ*_*X*_ is the slope factor. The T-type current inactivation variable (*b*), however, is described as follows:

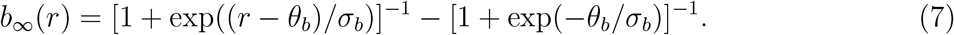

The calcium-activated, voltage-independent afterhyperpolarisation (AHP) potassium current (*I*_AHP_) is defined as follows:

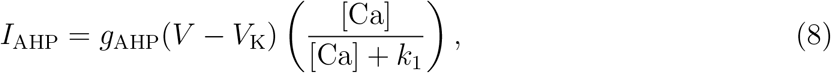

where *g*_AHP_ is the maximal conductance and *k*_1_ is the dissociation constant of the calciumdependent AHP current. The the intracellular concentration of calcium ions ([Ca]) is described by the following first-order differential equation:

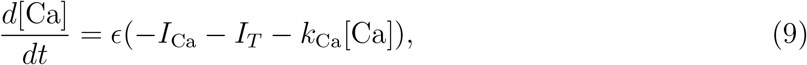

where ∈ is a constant that describes the effects of buffers, cell volume and the molar charge of calcium, and *k*_Ca_ denotes the calcium pump rate constant.

### 2.3 Synapse model

In Eq. (1), *I*_syn_ represents the synaptic current. For the STN cells, the synaptic current is only composed of the GPe’s contribution due to the lack of intrinsic connectivity within the STN, i.e., 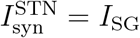. For the GPe cells, the synaptic current comprises two terms, i.e., 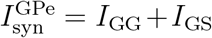, where *I*_GG_ (*I*_GS_) indicates GPe → GPe (STN → GPe) synaptic current:

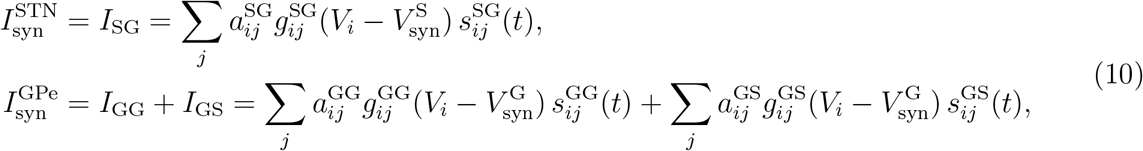

where 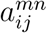, with *m, n* ∈ {STN (S), GPe (G)}, is the corresponding array of the adjacency ma-trix and 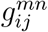 is the synaptic strength from the presynaptic neuron *j* in population *n* to the postsynaptic neuron *i* in population *m*. The parameter *V*_*i*_ is the membrane potential of the postsynaptic neuron and *V*_syn_ represents the corresponding synaptic reversal potential. The synaptic variable *s*_*ij*_(*t*) obeys the following first-order kinetics:

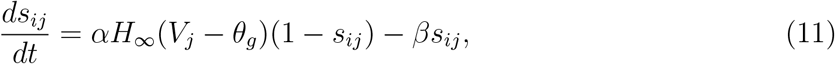

where *V*_*j*_ is the membrane potential of the presynaptic neuron, *α* and *β* are the opening and closing rates of channels, respectively, and *H*_∞_ is given by:

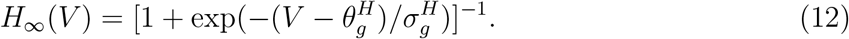

The numerical values of model parameters are presented in Tables 2 and 3.

**Table 2:**
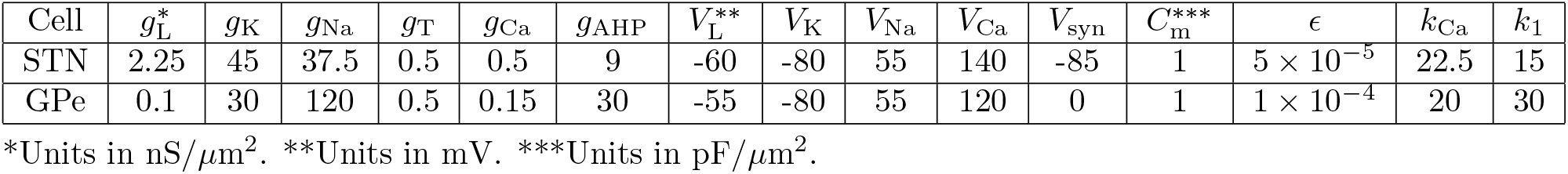
Parameters of maximal conductance (*g*_X_), neuronal and synaptic reversal potentials (*V*_X_), and calcium dynamics for the STN and GPe neuron models [11].

| Cell | $g_L^*$ | $g_K$ | $g_{Na}$ | $g_T$ | $g_{Ca}$ | $g_{AHP}$ | $V_L^{**}$ | $V_K$ | $V_{Na}$ | $V_{Ca}$ | $V_{syn}$ | $C_m^{***}$ | $\epsilon$ | $k_{Ca}$ | $k_1$ |
| --- | --- | --- | --- | --- | --- | --- | --- | --- | --- | --- | --- | --- | --- | --- | --- |
| STN | 2.25 | 45 | 37.5 | 0.5 | 0.5 | 9 | -60 | -80 | 55 | 140 | -85 | 1 | $5 \times 10^{-5}$ | 22.5 | 15 |
| GPe | 0.1 | 30 | 120 | 0.5 | 0.15 | 30 | -55 | -80 | 55 | 120 | 0 | 1 | $1 \times 10^{-4}$ | 20 | 30 |

\*Units in nS/ $\mu m^2$ . \*\*Units in mV. \*\*\*Units in pF/ $\mu m^2$ .

**Table 3:** Kinetic parameters for the STN and GPe neuron models [11].

| Cell | $X$ | $\theta_X$ | $\sigma_X$ | $\tau_X^{0*}$ | $\tau_X^1$ | $\theta_X^\tau$ | $\sigma_X^\tau$ | $\phi_X$ |
| --- | --- | --- | --- | --- | --- | --- | --- | --- |
| STN | $n$ | -32 | 8 | 1 | 100 | -80 | -26 | 0.75 |
| | $m$ | -30 | 15 | — | — | — | — | — |
| | $h$ | -39 | -3.1 | 1 | 500 | -57 | -3 | 0.75 |
| | $a$ | -63 | 7.8 | — | — | — | — | — |
| | $r$ | -67 | -2 | 40 | 17.5 | 68 | -2.2 | 0.2 |
| | $s$ | -39 | 8 | — | — | — | — | — |
| | — | $\theta_g$ | $\theta_g^H$ | $\sigma_g^H$ | $\alpha$ | $\beta$ | $\theta_b$ | $\sigma_b$ |
|  | — | 30 | -39 | 8 | 5 | 1 | 0.4 | -0.1 |
| GPe | $n$ | -50 | 14 | 0.05 | 0.27 | -40 | -12 | 0.05 |
| | $m$ | -37 | 10 | — | — | — | — | — |
| | $h$ | -58 | -12 | 0.05 | 0.27 | -40 | -12 | 0.05 |
| | $a$ | -57 | 2 | — | — | — | — | — |
| | $r$ | -70 | -2 | 30 | — | — | — | 1 |
| | $s$ | -35 | 2 | — | — | — | — | — |
| | — | $\theta_g$ | $\theta_g^H$ | $\sigma_g^H$ | $\alpha^{**}$ | $\beta^{**}$ | — | — |
|  | — | 20 | -57 | 2 | 2 | 0.08 | — | — |

\*Units in ms. \*\*Units in $ms^{-1}$ .

### 2.4 Rhythmic stimulation paradigm

In Eq. (1), *I*_stim_(*t*) represents the stimulation current composed of continuous cathodic pulses with a specific amplitude and frequency (also see Fig. 2A), that conceptually resembled the waveform of DBS [64]:

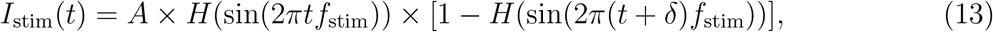

where *A* is the amplitude, *f*_stim_ = 1/*T*_stim_ is the frequency of the stimulation signal, and *T*_stim_ denotes the period of stimulation (i.e., inter-pulse interval). The parameter *δ* represents the pulse width (i.e., duration of each stimulus), and *H*(*x*) is the Heaviside step function, such that *H*(*x*) = 0 if *x* ≤ 0, and *H*(*x*) = 1, otherwise.

**Figure 2:**
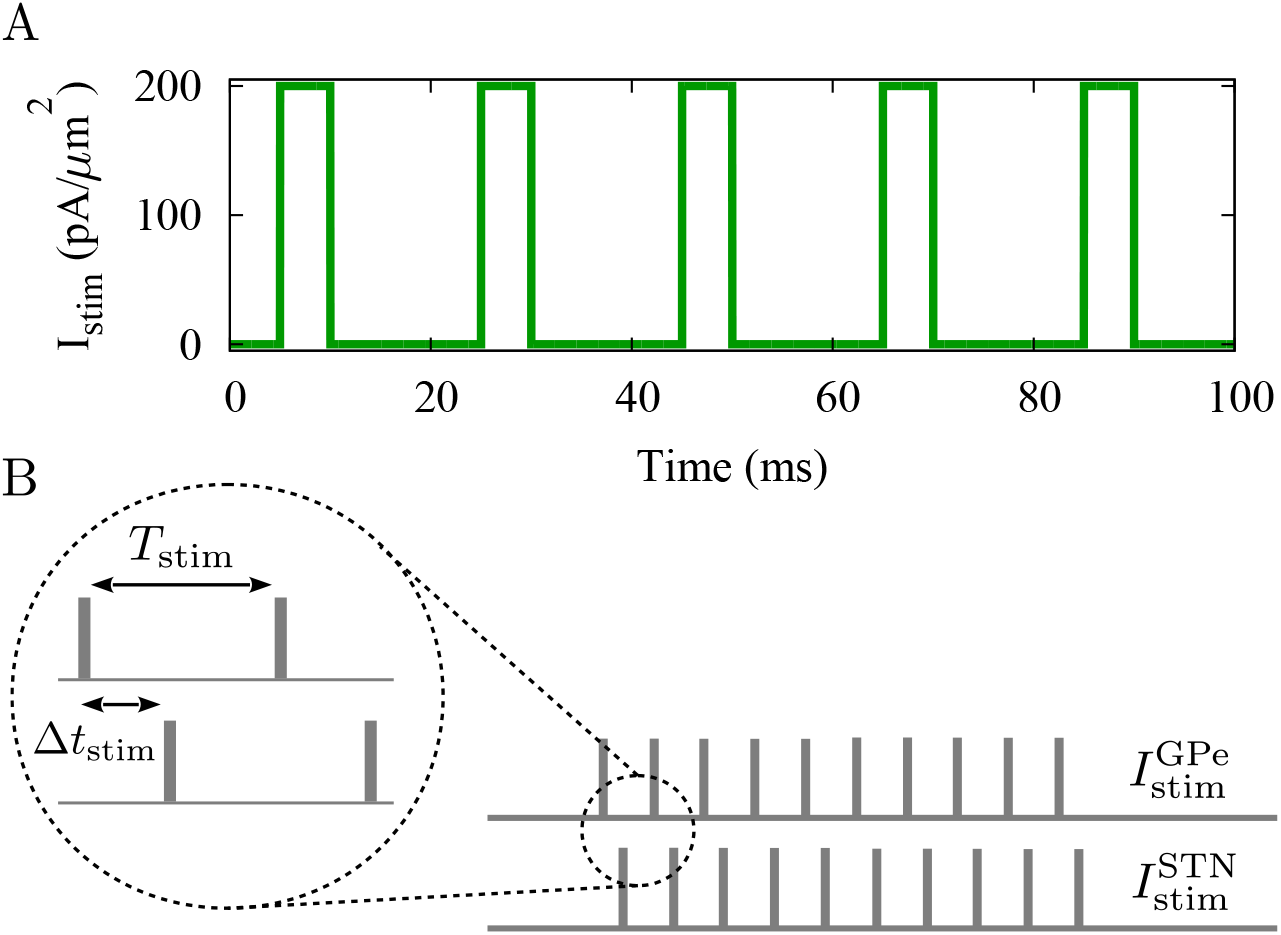
The rhythmic stimulation paradigm. (**A**) Representative time course of a simulated stimulation current generated by Eq. (13), with *A* = 200 pA/*µ*m^2^, *f*_stim_ = 50 Hz and *δ* = 5.0 ms. (**B**) Schematics of the stimulation signals delivered to the STN 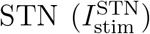and GPe 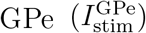, each composed of a series of cathodic rectangular pulses. The symbol *T*_stim_ represents the period of stimulation or inverse frequency (1/*f*_stim_) and Δ*t*_stim_ denotes the time shift between the two stimulation signals delivered to the STN and GPe.

The stimulation was assumed to be global, i.e., the stimulation signal was simultaneously delivered to all cells within the target population for a specific duration (i.e., the *stimulation epoch*), uniformly affecting all cells. Furthermore, as it is shown in Fig. 2B, a crucial aspect of the rhythmic stimulation paradigm in our model is the introduction of a time shift (Δ*t*_stim_) [30] between the two stimulation signals that are separately delivered to the two target populations (i.e., STN and GPe). This time shift represents the delay between the onset times of pulses within the respective stimulation currents delivered to the two populations, allowing for targeting plasticity.

### 2.5 Inhibitory spike-timing-dependent plasticity (iSTDP)

The GPe→ STN synaptic connections characterised by the strength *g*_SG_ were subjected to the following symmetric iSTDP profile [39, 41]:

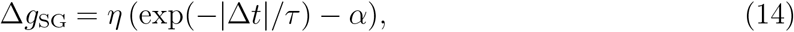

where *η* is the learning rate, *τ* is the decay time constant of the exponential function and *α* is the depression factor, as given in Table 4 (used as the *default iSTDP profile* throughout the manuscript). The time lag Δ*t* = *t*_post_−*t*_pre_ represents the instantaneous time difference between pre- and postsynaptic spike pairs. The synaptic strengths were updated by an additive rule at each step of the simulation, i.e., *g* → *g* + Δ*g*.

**Table 4:** The iSTDP model parameters [66].

| Parameter | Symbol | Value |
| --- | --- | --- |
| Learning rate | $\eta^*$ | 0.005 |
| Decay time constant | $\tau^{**}$ | 20 |
| Depression factor | $\alpha$ | 0.2 |

\*Units in nS/ $\mu\text{m}^2$ . \*\*Units in ms.

The synaptic strengths were confined in the range [*g*_min_, *g*_max_] ∈ [0.05, 0.5] nS/*µ*m^2^, and were set to *g*_min_ (*g*_max_) by using a hard bound saturation constraint once they crossed the lower (upper) bound of their allowed range. A non-zero lower bound (*g*_min_) guarantees the stabilisation of the network synaptic connectivity in a loosely connected state [44].

### 2.6 Analysis of network activity and connectivity

#### 2.6.1 Mean firing rate

The firing rate of individual neurons was estimated as the average spike count over the entire simulation period, excluding the 500 ms of initial network transients. The mean network firing rate was then derived by averaging the firing rates of all neurons in the network.

#### 2.6.2 Local field potential (LFP)

The rhythmic activity of the STN and GPe is reflected in well-pronounced oscillations of their simulated local field potential (LFP) which is an indicator of synchronised neuronal dynamics. The LFP was defined as the weighted sum of the membrane potential of all neurons in a population, as follows [65]:

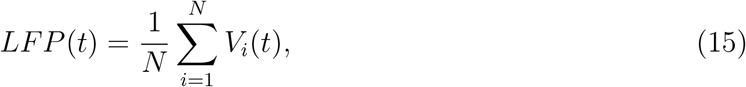

where *V* (*t*) is the membrane potential whose evoultion is given by Eq. (1), and *N* represents the total number of neurons in each population.

#### 2.6.3 Synchrony index

The global synchrony of the STN or GPe was quantified by calculating the neuronal coherency (*χ*_*p*_) within the population *p*, as follows [54, 67]:

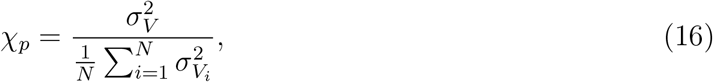

where 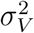 is the time-averaged variance of the global membrane potential of all neurons in a pop-ulation, and 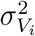is the variance of membrane potential of the *i*-th neuron. This measure quantifies the average fluctuations of the neuronal membrane potentials to infer synchrony, normalised in the range [0, 1], i.e., *χ*_*p*_ = 0 represents an incoherent state when the fluctuations of membrane potential are zero and *χ*_*p*_ = 1 when all the neurons have the same trajectory [67].

For synchronous activity in the STN-GPe network, synchrony must be be present in both the STN and GPe. Therefore, synchrony in the STN-GPe network (*χ*) was estimated as the geometric mean of the individual synchrony of each population [54]:

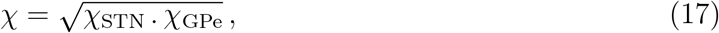

which is calculated once the system has converged to a stable state. In this setting, synchrony reflects the correlated neuronal activity in the STN-GPe network.

#### 2.6.4 Phase-locking analysis

To quantify the spike timing synchronisation relative to the rhythmic stimulation waveform, the phase-locking value (PLV) was calculated as follows [68]:

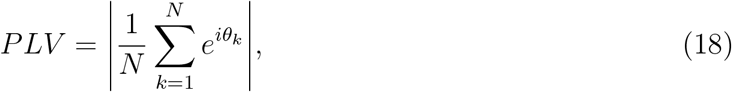

where *N* is the number of spikes and *θ*_*k*_ is the phase of the stimulation at which the *k*-th spike occurs. A uniform distribution of spike timings over all phases results in a PLV of 0, i.e. no synchronisation, while a PLV value of 1 indicates perfect synchronisation.

#### 2.6.5 Power spectral density (PSD)

To estimate the strength of oscillations, the power spectral density (PSD) of the collective activity in each population was calculated by the fast Fourier transform (FFT) of the oscillatory signal with a sampling frequency of 50 kHz.

#### 2.6.6 Weighted connectivity and synaptic wiring

To find the final network topology based on the GPe→ STN synaptic strengths, the structural connectivity matrix (i.e., the adjacency matrix) was transformed into a binary weighted connectivity matrix. This was performed by introducing a threshold (*h*) over which the link between two neurons is maintained [30], i.e., weighted connectivity matrix array, *c*_*ij*_ = 1 if *g*_*ij*_ ≥ *h*, and *c*_*ij*_ = 0, otherwise. The synaptic wiring was then calculated by counting the number of links pre- and post-stimulation compared to the structural connectivity, which was set as a reference value (100% wiring) at the initial setting. For simplicity, wiring is reported by a number in the range [0, 1] such that zero wiring represents a disconnected network, whereas wiring close to unity indicates a fully connected network.

## 3 Results

### 3.1 The STN-GPe network dynamics

The structural characteristics of the STN-GPe network are shown in Fig. 3A, where neurons in the GPe→ GPe, GPe→ STN, and STN→ GPe pathways are randomly connected according to the connection probabilities given in Table 1. The GPe → STN connectivity strengths were selected from a normal distribution whose mean was set at the midpoint of the allowed range, ensuring an unbiased initial configuration (see Fig. 3B). The GPe → GPe and STN→ GPe synaptic strengths were set similarly. The neuronal and network activity patterns pertaining to healthy and PD conditions were realised by adjusting the external currents applied to the STN and GPe (also see Table 5) [11, 42]. This was motivated by experimental findings in rodents suggesting that striato-pallidal inhibition [69] and cortico-subthalamic excitation [70] are augmented following dopamine loss. In the model, the external currents applied to the STN and GPe were selected from normal distributions designed to reflect healthy and PD states (Fig. 3C).

**Table 5:**
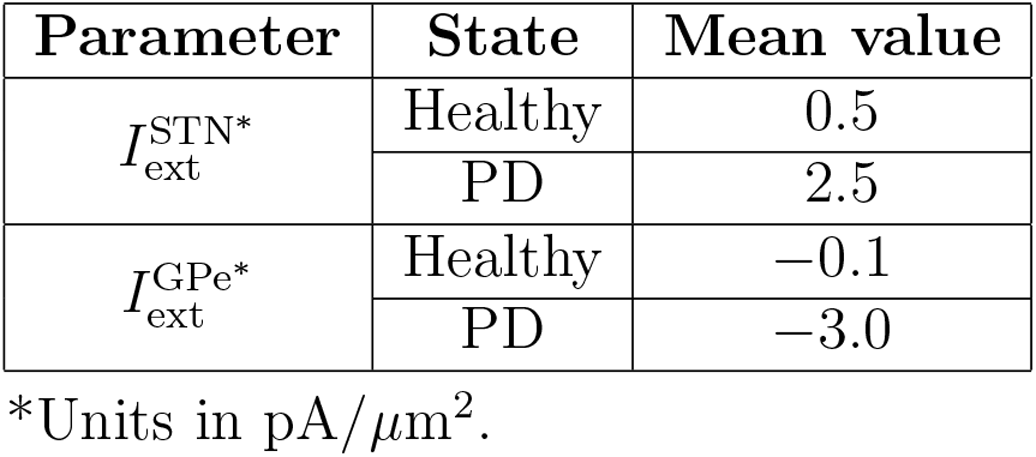
External currents in control and PD states [42].ss.

**Figure 3:**
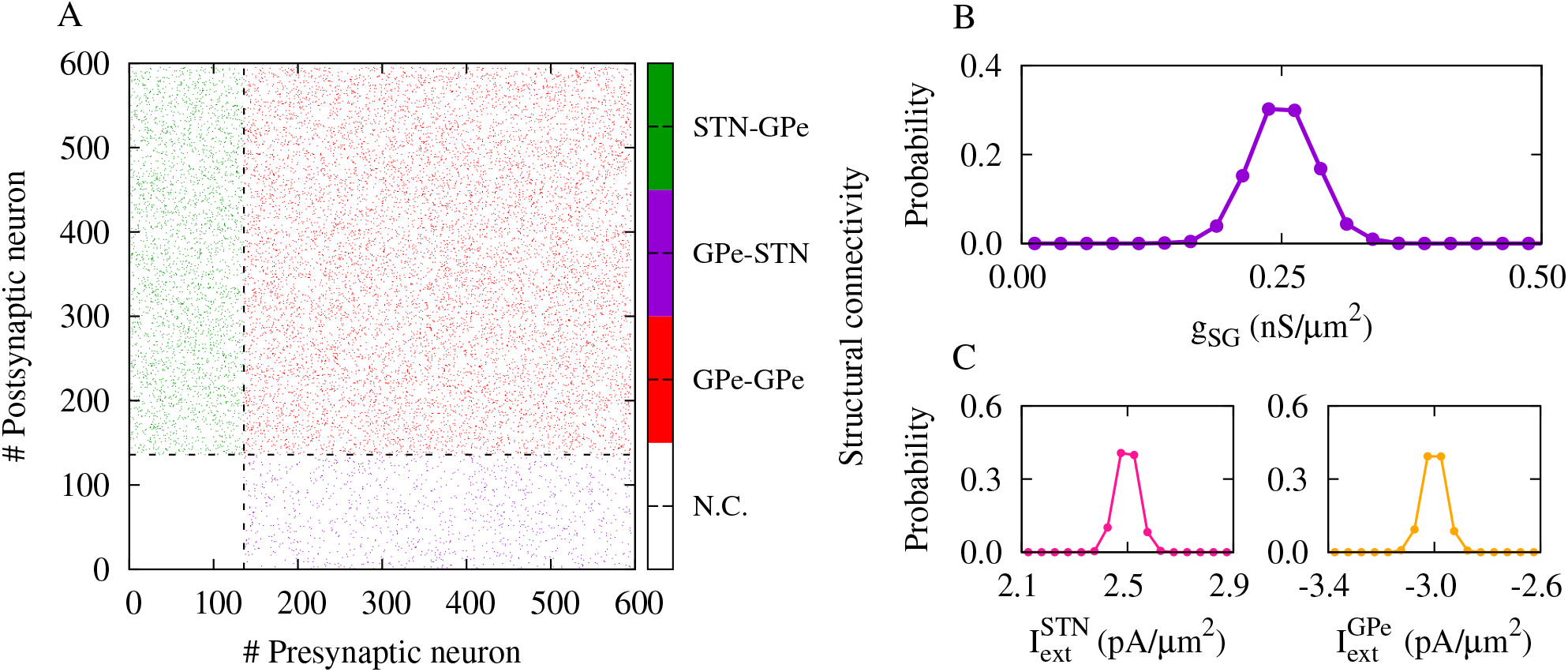
Network topology and model inputs. (**A**) Structural connectivity of the network calculated using a binary representation, i.e., adjacency matrix array, *a*_*ij*_ = 1 (coded by colour) if the two neurons are connected, and *a*_*ij*_ = 0 (white; N.C. stands for *not connected*), otherwise. Dashed lines indicate the STN (# 1-136) and GPe (# 137-460) neurons. (**B**) Normal distribution of the (magnitude of) GPe→ STN synaptic strengths around 0.25 nS/*µ*m^2^. (**C**) Normal distribution of the external currents applied to the STN (left) and GPe (right) in the PD state around 2.5 pA/*µ*m^2^ and −3.0 pA/*µ*m^2^, respectively.

The oscillatory dynamics of the STN (pink) and GPe (orange) in healthy (Fig. 4A1) and PD (Fig. 4A2) states are reflected in their respective rater plots (upper panels) and LFPs (lower panels). The model parameters including external currents and synaptic strengths were tuned to obtain mean firing rates (Fig. 4B) in the STN (healthy: ∼15 Hz, and PD: ∼10 Hz) and GPe (healthy: ∼35 Hz, and PD:∼ 23 Hz) in the experimentally observed ranges in healthy and parkinsonian rodents [8, 9, 71]. More specifically, in the healthy state the activity of the STN-GPe network is weakly synchronised (Fig. 4C, blue) as evidenced by small-amplitude oscillations in the LFP (Fig. 4A1), and sharp peaks are absent in the PSD of population activity of each nuclei in the beta (15-30 Hz) frequency range (Fig. 4D, left). In the PD state, however, the mean firing rate of the STN (GPe) is increased (decreased) compared to the healthy state (cf. Fig. 4B, blue and red). In this case, the activity of the STN-GPe network is strongly synchronised (Fig. 4C, red) as reflected in well-pronounced LFP oscillations (Fig. 4A2), and beta frequency peaks are present in the PSD of population activity of each nuclei where the power of beta oscillations is also increased (Fig. 4D, right).

**Figure 4:**
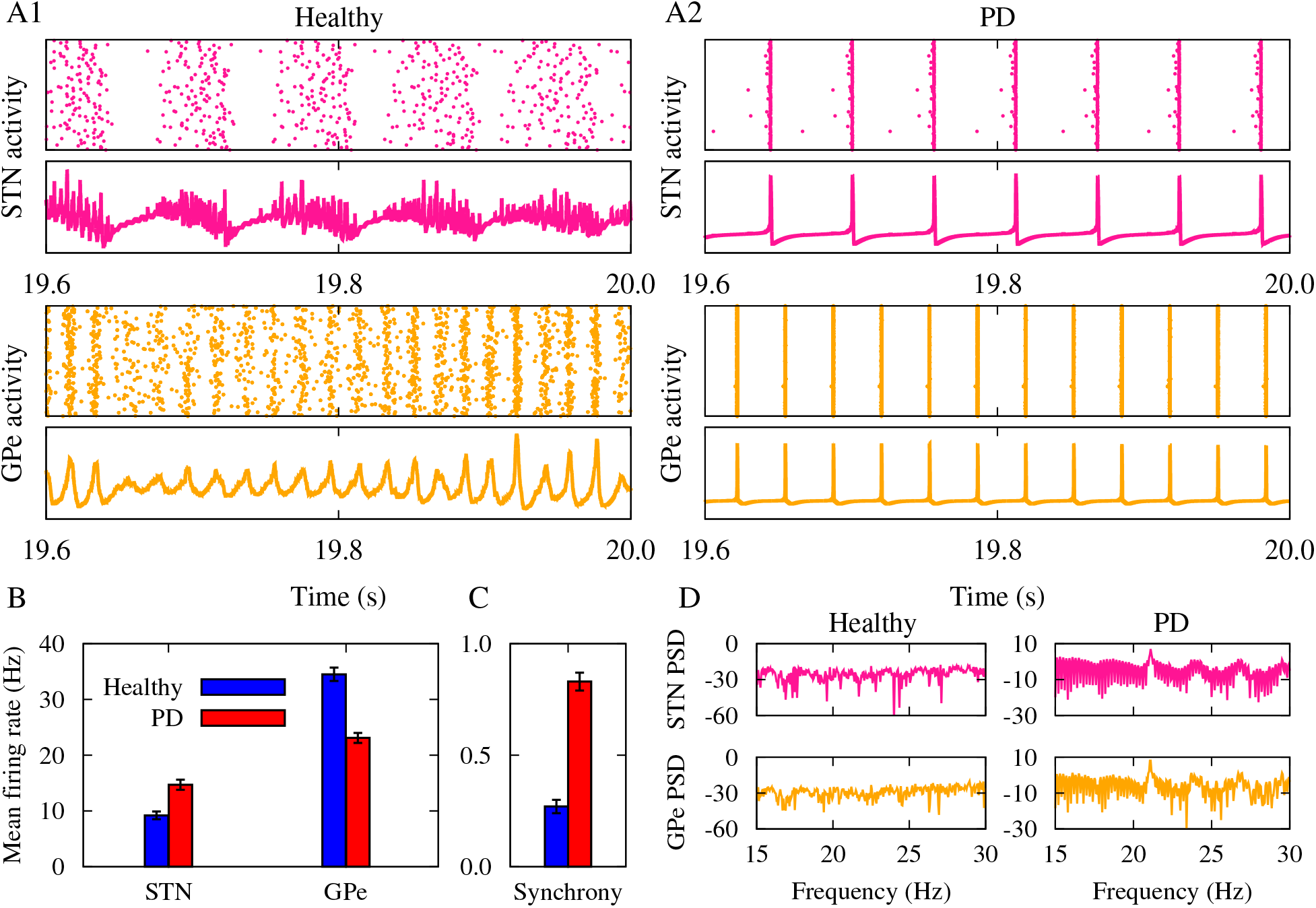
Oscillatory dynamics of the STN-GPe network. (**A1, A2**) Raster plot (upper panel) and LFP (lower panel) of the STN and GPe in healthy (A1) and PD (A2) states. (**B**) Mean firing rate of the STN and GPe in healthy (blue) and PD (red) states. (**C**) The STN-GPe synchrony in healthy (blue) and PD (red) conditions. (**D**) PSD of the STN and GPe activity in healthy (left) and PD (right) states.

### 3.2 Theoretical prediction of synaptic rewiring

To investigate how a plastic synaptic wiring pattern may influence STN-GPe oscillations, in the model we assumed that the GPe→ STN synaptic strengths are modified based on a generic, temporally symmetric iSTDP profile [39, 41] given by Eq. (14), as shown in Fig. 5A. According to the iSTDP rule, the synapse is strengthened (Fig. 5A, red region) whenever the difference between the spike timings of neuron pairs is smaller than a critical value, i.e., |Δ*t*^∗^| = −*τ* ln*α* based on Eq. (14), whereas the synapse is weakened otherwise (Fig. 5A, blue region). Of note, this differs from the temporally asymmetric classical STDP rule of excitatory synapses [72–74], where pre-post or post-pre pairing of spikes determines the nature of the synaptic change, i.e., long-term potentiation or long-term depression, respectively.

**Figure 5:**
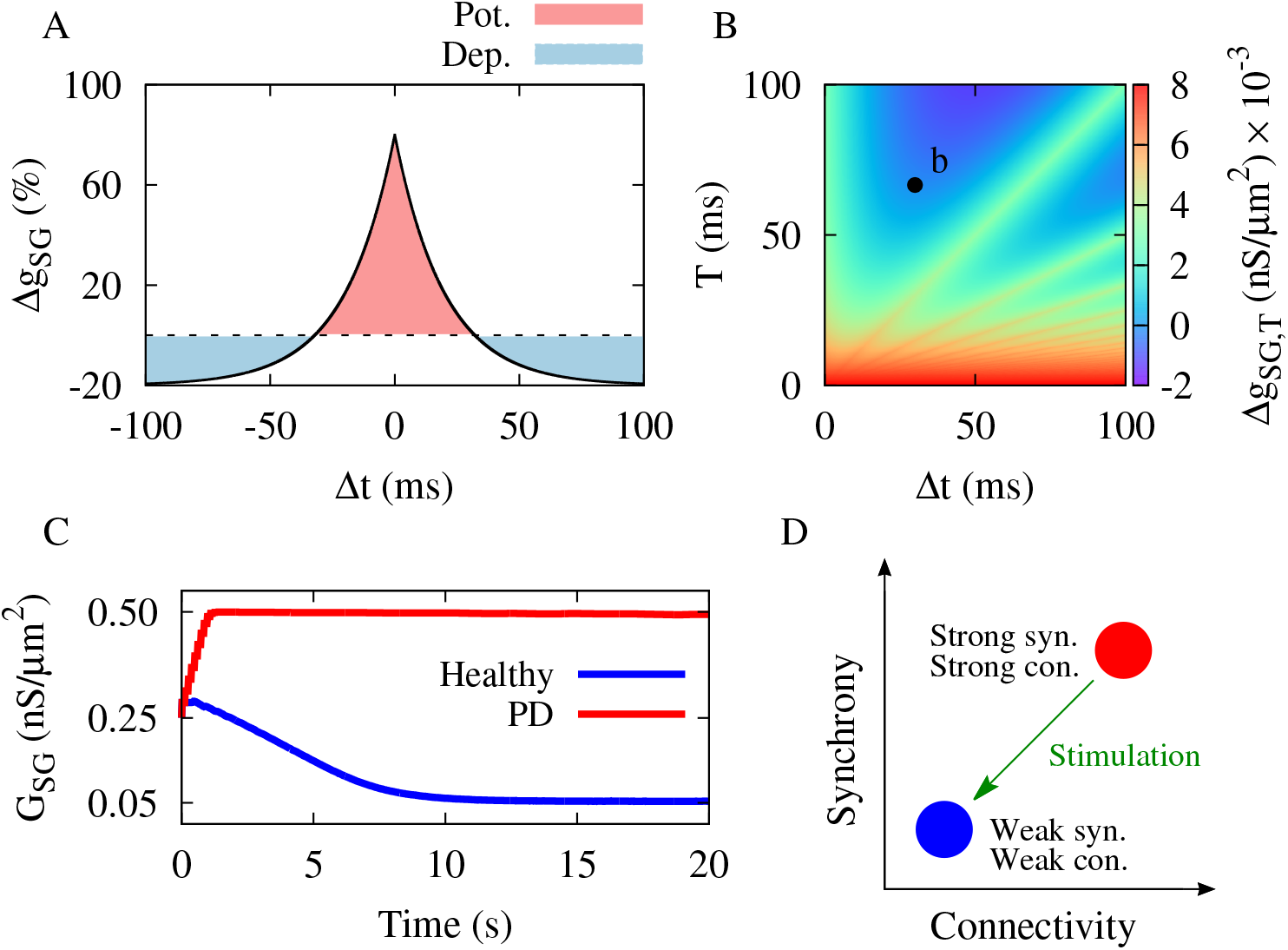
Synaptic rewiring by iSTDP. (**A**) Synaptic change due to the temporally symmetric iSTDP profile described by Eq. (14) with parameters given in Table 3. Red (blue) region indicates synaptic potentiation (depression). Dashed line indicates Δ*g*_SG_ = 0. (**B**) Theoretical prediction of the colour-coded synaptic change based on Eq. (19) with red (blue) representing synaptic potentiation (depression). Point b with (Δ*t, T*) = (30, 66.66) indicates the default stimulation setting. (**C**) Numerical simulation of the down- (blue) and up-regulation (red) of the mean GPe→ STN synaptic coupling in healthy (blue) and PD (red) states. (**D**) Schematics of a bistability between healthy (blue; weak synchrony and weak connectivity) and PD states (red; strong synchrony and strong connectivity). External stimulation (green) shifts network dynamics from the PD state to the healthy state.

When evaluated over the spiking cycle (*T*), a synapse shared by two neurons experiences a succession of two weight modifications (Δ*g*_T_) characterised by the two time lags, i.e., Δ*t* and *T* − Δ*t*, as follows [30]:

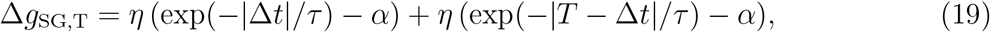

where the net synaptic change over the spiking cycle is determined by the sum of the two terms, i.e., one for synaptic potentiation and the other for synaptic depression.

Fig. 5B represents a theoretical prediction of synaptic rewiring by estimating the amount of synaptic change based on Eq. (19) when the time lag between spike pairs (Δ*t*) and the spiking cycle (*T*) are systematically varied as free parameters. The colour-coded synaptic change indicates that for small values of the spiking cycle (*T* < 20 ms) the synapses undergo strong potentiation (red). In contrast, for higher values of the spiking cycle (*T* > 20 ms), the synapses are more likely to experience small potentiation (green) or even depression (blue) in the range*T* > 50 ms and 20 < Δ*t* < 80 ms.

Numerical simulations of the STN-GPe network confirmed theoretically predicted down-(blue) and up-regulation (red) of the mean GPe→ STN coupling by iSTDP in healthy and PD states, respectively, as shown in Fig. 5C. Together, the results presented in Figs. 4 and 5 suggest that both synchrony and connectivity in the STN-GPe network are modulated in healthy and PD states. In fact, in the healthy state, uncorrelated activity in the STN and GPe leads to the weakening of the GPe → STN synaptic connections. In contrast, in the PD state, synchronous STN-GPe activity strengthens the GPe → STN synapses through iSTDP (see Fig. 5C).

This is in line with previous results indicating that synaptic plasticity can shape qualitatively different attractor states in a variety of neuronal network models characterised by different regimes of activity and connectivity [42–45]. In our model, this relates to a bistability between the healthy state (Fig. 5D, blue ball) characterised by weak network synchrony (Fig. 4C, blue) and weak connectivity (Fig. 5C, blue) and the PD state (Fig. 5D, red ball) characterised by strong network synchrony (Fig. 4c, red) and strong connectivity (Fig. 5c, red).

### 3.3 Synaptic rewiring and modulation of oscillatory activity by time-shifted stimulation

As shown computationally [27], stimulation (Fig. 5D, green) can shift network dynamics from pathological states (Fig. 5D, red ball) to more physiologically favoured states (Fig. 5D, blue ball). Here, we show that stimulation-induced network state transitions are mediated by synaptic rewiring of the network, as well as changes in network activity. We demonstrate that a specific pattern of stimulation with an appropriate time shift (Δ*t*_stim_) and frequency (*f*_stim_ = 1/*T*_stim_) allows us to harness plasticity mechanisms to drive favourable changes in the synaptic strengths and, thereby, effectively shift the dynamics of the STN-GPe network from the PD state to the healthy state and induce an effect that outlasts the cessation of stimulation.

To specifically target plasticity for synaptic rewiring, we chose stimulation time shift and frequency from theoretically predicted ranges of parameters in Fig. 5B, which were favourable for synaptic weakening (blue regions), i.e., point b with Δ*t*_stim_ = 30 ms and *f*_stim_ = 15 Hz (*T*_stim_= 66.66 ms) as the *default stimulation setting* in the manuscript. To be able to selectively modulate the inter-population synaptic connections, we first tested whether the stimulation could force neurons to spike whenever applied to each population. In principle, this allows for a prolonged synaptic rewiring by paired simulation-induced Hebbian plasticity between the two inter-connected populations, as demonstrated both experimentally and computationally [55,56]. Raster plots in Fig. 6A show that the rhythmic stimulation forces neuronal spikes in the STN and GPe upon administration, where the phases of individual neurons in each population are locked to the stimulation waveform as assessed by PLV for each population in Fig. 6B.

**Figure 6:**
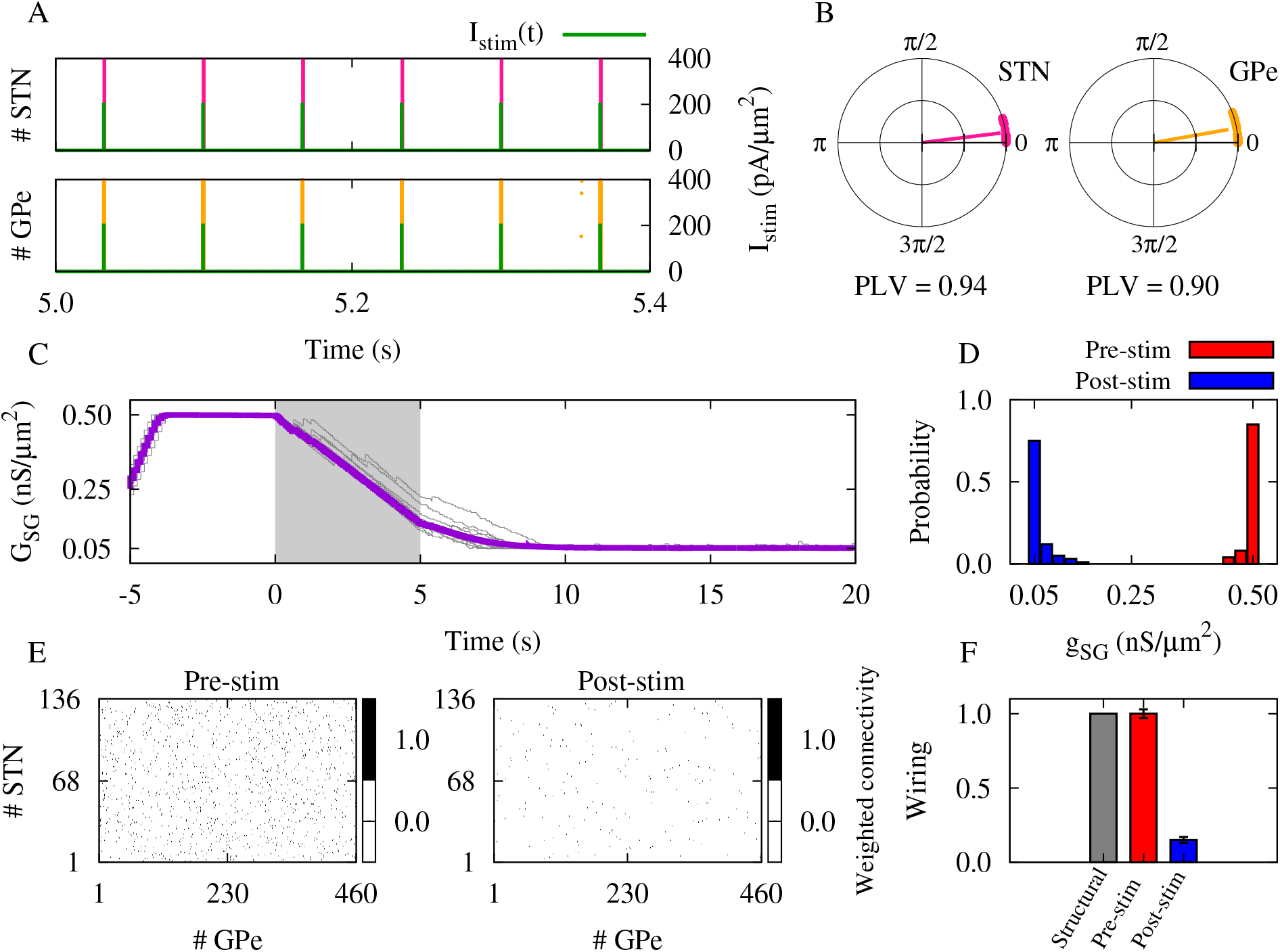
Synaptic rewiring by dual STN-GPe stimulation. (**A**) Raster plot of the STN (top) and GPe (bottom) activity locked to the rhythmic stimulation (green). (**B**) PLV (the radial bar) and polar distribution of the phases of the STN (left) and GPe (right) neurons in the presence of stimuli. (**C**) Time course of the mean GPe→ STN synaptic coupling (violet) and ten randomly chosen individual synaptic strengths (gray curves). The shaded area indicates the stimulation epoch (0 < *t* < 5 s). (**D**) Pre- (red) and post-stimulation (blue) distribution of the GPe→ STN synaptic strengths. (**E**) Binary representation of the weighted GPe→ STN connectivity matrix constructed by introducing a threshold (*h* = 0.1) pre- (left) and post-stimulation (right). (**F**) Pre- (red) and post-stimulation (blue) GPe →STN synaptic wiring with respect to the structural connectivity (gray) as a reference.

Then, we tuned the STN-GPe network to model the PD state recognised by strong network synchrony and strong GPe→ STN synaptic weights (Fig. 6C, *t* < 0 s). The rhythmic stimulation was turned ON at *t* = 0 s, uniformly affecting all cells within the STN and GPe for the stimulation epoch of 5 s (Fig. 6C, shaded area), and then was turned OFF. The evolution of synaptic strengths in Fig. 6C shows that strong GPe→ STN synaptic connections before stimulation (*t* < 0 s) are down-regulated during the stimulation epoch (0 < *t* < 5 s), and this weakening of the synaptic strengths persists after cessation of the stimulation (*t* > 5 s), as reflected in the mean synaptic weight (violet) as well as in randomly chosen individual synaptic strengths (gray). Consequently, the synaptic strengths are redistributed from an initially strong distribution pre-stimulation (Fig. 6D, red) to a finally weak setting post-stimulation (Fig. 6D, blue). As a result, the weighted connectivity (see Methods) switches from an increased connectivity state pre-stimulation to a decreased connectivity state post-stimulation (Fig. 6E), as demonstrated by the reduced post-stimulation wiring (∼ 0.15) compared to the pre-stimulation wiring (∼ 0.95) as well as the structural wiring (= 1) as a reference (Fig. 6F).

Notably, the reduction in synaptic weights during the stimulation epoch (0 < *t* < 5 s) and even after the cessation of stimulation (*t* > 5 s) points out the potential transnational implications of this stimulation method by inducing long-lasting after-effects beyond the stimulation epoch. In this context, the continuation of synaptic weakening is a feedback process mediated by the iSTDP profile (Fig. 5A). Prior to stimulation, highly correlated, synchronised activity of the STN and GPe neurons potentiates the inter-population synaptic strengths. Administration of rhythmic stimulation with a specific time shift and frequency relative to endogenous spiking activity disrupts spike timings and forces neurons to fire at a rate designed to induce synaptic weakening. As shown previously [30], sufficient down-regulation of the synaptic strengths be-low a certain threshold during the stimulation epoch prevents the re-potentiation of synaptic strengths after the cessation of stimulation. Together, these changes stabilise synaptic connectivity in a loosely connected state (see Fig. 6E and F).

Interestingly, the oscillatory activity of STN-GPe is modulated following synaptic rewiring by time-shifted stimulation, as demonstrated in Fig. 7. The mean firing rates of the STN and GPe are reset to their healthy ranges (cf. Fig. 7A and Fig. 4B). Stimulation-induced reduction of the synaptic strengths ultimately desynchronises the STN-GPe activity post-stimulation (Fig. 7B, blue) compared to highly synchronised activity pre-stimulation (Fig. 7B, red). In addition, the PSD of the STN and GPe activity shows a reduction of the beta oscillatory power post-stimulation (Fig. 7C, right) compared to increased beta activity pre-stimulation (Fig. 7C, left), suggesting that suppression of abnormal beta activity is feasible by targeted reshaping of synaptic connectivity.

**Figure 7:**
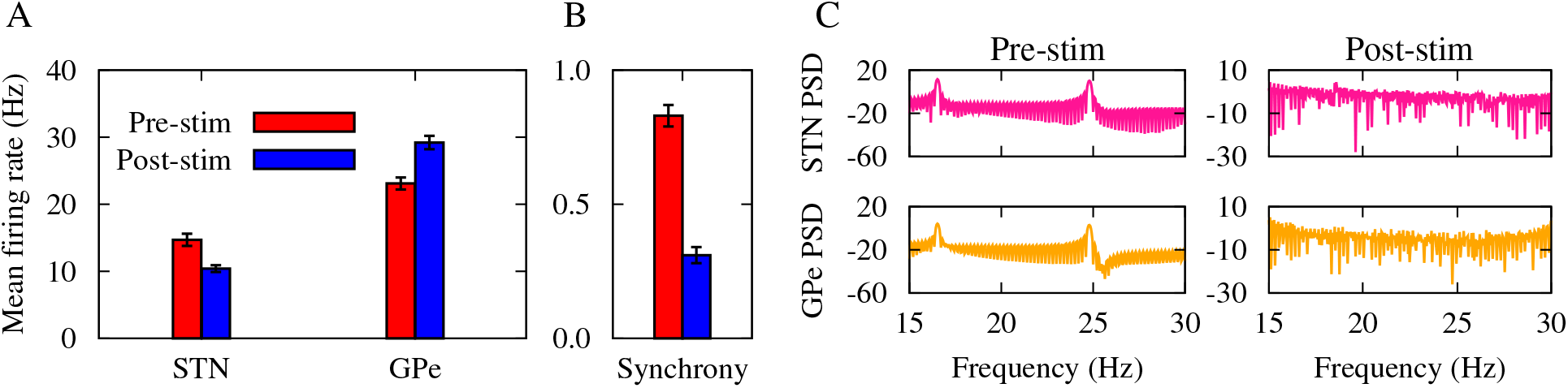
Modulation of oscillatory activity following synaptic rewiring. (**A**) Mean firing rate of the STN and GPe pre- (red) and post-stimulation (blue). (**B**) The STN-GPe synchrony pre- (red) and post-stimulation (blue). (**C**) PSD of the STN and GPe activity pre- (left) and post-stimulation (right).

### 3.4 Temporal pattern of stimulation

In standard DBS, stimuli are delivered continuously at a constant rate. However, to reduce the integral amount of current administered, temporally patterned stimulation protocols were developed and tested in PD patients and pre-clinical models [75]. To demonstrate whether mutually shifting pulse patterns by Δ*t*_stim_ works in a generic way and to explore the influence of temporal intermittency in the stimulation signal on post-stimulation STN-GPe synchrony and GPe→STN synaptic wiring, we repeated our simulations with burst stimulation by utilising bursts of stimuli (illustrated in Fig. 8A), as follows [30]:

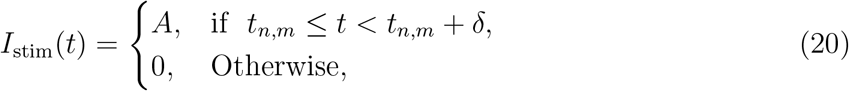

where *t*_*n,m*_ = [1000*n*/*f*_stim_ + *m*(*T*_ON_ + *T*_OFF_)] ms with *n* = 0, 1, 2, … are the times of the pulse onsets, *m* is the burst counter with *b* = 5 pulses within each burst, *f*_stim_ = 15 Hz is the (intraburst) frequency of stimulation, *T*_ON_ = [1000(*b* − 1)/*f*_stim_ + *δb*] ms is the duration of each burst sand *T*_OFF_ = 500 ms is the inter-burst interval (see Fig. 8A).

**Figure 8:**
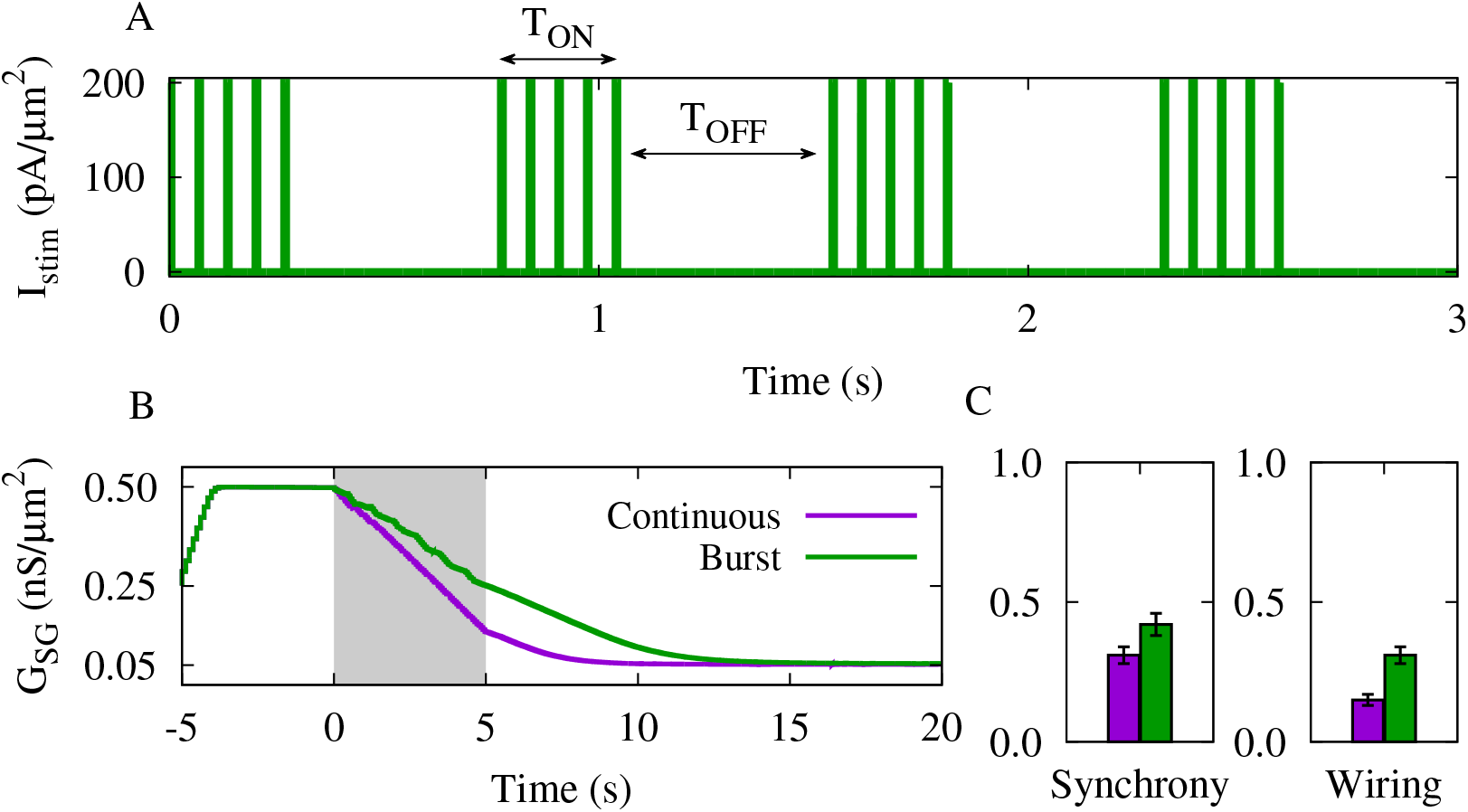
The effect of intermittent stimulation on synchrony and synaptic rewiring. The outcome of stimulation is compared for the continuous (violet) and burst (green) stimuli.(**A**) Time course of the burst stimulation generated by Eq. (20), with *A* = 200 pA/*µ*m^2^, *f*_stim_ = 15 Hz, *T*_OFF_ = 500 ms and *δ* = 5.0 ms. (**B**) Time course of the mean GPe→ STN synapticcoupling. The shaded area indicates the stimulation epoch (0 < *t* < 5 s). (**C**) The STN-GPe synchrony (left) and GPe → STN synaptic wiring (right).

Burst stimulation was administered with the same intra-burst frequency as the continuous stimulation (i.e., *f*_stim_ = 15 Hz), as shown in Fig. 8A. Similarly to the continuous stimulation scenario, the two stimulation signals were delivered to the STN and GPe with a time shift Δ*t*_stim_ = 30 ms. The time course of the mean GPe→ STN synaptic coupling in Fig. 8B demonstrates that during the stimulation epoch (shaded area) of the burst stimulation (green) the reduction of the synaptic weights occurs in a step-like manner due to the ON/OFF cycles (see Fig. 8A). This ultimately relaxes the synaptic strengths during the *T*_OFF_ cycle, preventing a monotonic reduction of the synaptic strengths during the stimulation epoch. Therefore, as shown in Fig. 8C the continuous stimulation (violet) comparatively outperforms the burst stimulation (green) in reducing the post-stimulation STN-GPe synchrony (left) and GPe→ STN synaptic wiring (right).

### 3.5 Charge-balanced stimulation waveform

Standard DBS and conceptually similar brain stimulation protocols often utilise charge-balanced stimuli to mitigate the risk of tissue damage [76]. Therefore, we repeated our simulations to test whether the outcome of the rhythmic stimulation depends on the use of charge-balanced stimuli. As shown in Fig. 9A, we used a symmetric biphasic stimulation waveform with zero inter-phase delay in our model to yield a short cathodic rectangular pulse followed by another pulse with the opposite polarity and the same duration, resulting in a zero net charge. To that end, the stimulation signal was implemented as follows [77]:

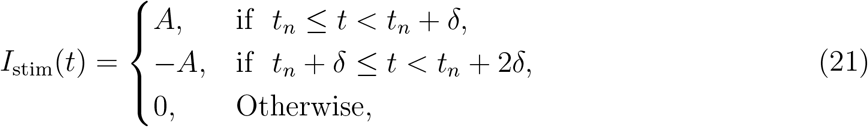

where *t*_*n,m*_ = [1000*n*/*f*_stim_] ms with *n* = 0, 1, 2, … are the times of the pulse onsets, and *f*_stim_ = 15 Hz is the frequency of stimulation.

**Figure 9:**
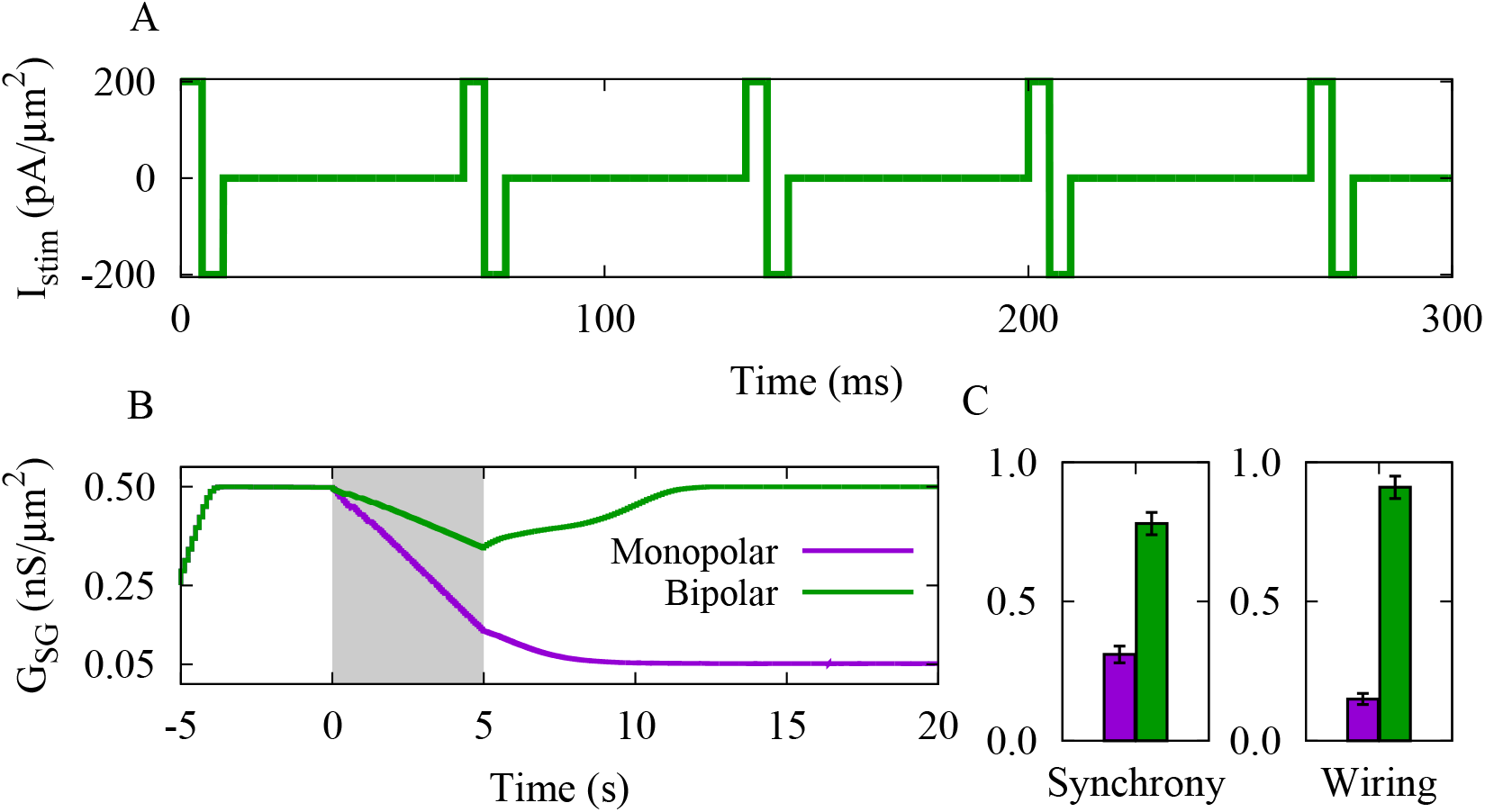
The effect of bipolar stimulation on synchrony and synaptic rewiring. The outcome of stimulation is compared for the monopolar (violet) and bipolar (green) stimuli.(**A**) Time course of the bipolar stimulation generated by Eq. (21), with *A* = *±*200 pA/*µ*m^2^,*f*_stim_ = 15 Hz and *δ* = 5.0 ms. (**B**) Time course of the mean GPe→ STN synaptic coupling.The shaded area indicates the stimulation epoch (0 < *t* < 5 s). (**C**) The STN-GPe synchrony (left) and GPe → STN synaptic wiring (right).

Bipolar stimulation was administered with the same frequency as the monopolar stimulation (i.e., *f*_stim_ = 15 Hz), as shown in Fig. 8A, with the same time shift Δ*t*_stim_ = 30 ms as the default stimulation setting. The mean GPe→ STN synaptic coupling in Fig. 9B shows that during the stimulation epoch the bipolar stimulation (green) reduces synaptic weights at a slower pace than the monopolar stimulation (violet). But, in this case, the synaptic strengths are not sufficiently reduced to prevent re-potentiation after stimulation offset, resulting in the growing of the synaptic weights post-stimulation. Consequently, as shown in Fig. 9C the post-stimulation STN-GPe synchrony (left) and GPe→ STN synaptic wiring (right) for bipolar stimulation (green) are not reduced to the same degree as for monopolar stimulation (violet). This can be resolved by an extended stimulation epoch (e.g., 10 s instead of 5 s).

### 3.6 Stimulation parameters and plasticity profile

Ultimately, the benefits of time-shifted stimulation may depend on its robustness against the re-tuning of stimulation parameters. So far, we used a predefined stimulation parameter set as the default stimulation setting, i.e., time shift Δ*t*_stim_ = 30 ms and frequency *f*_stim_ = 15 Hz. To test the effectiveness of the stimulation concept (i.e., time-shifted dual-site stimulation) in other ranges, we changed the stimulation time shift and frequency and observed the resultant change in the STN-GPe synchrony (Fig. 10A1) and synaptic wiring (Fig. 10A2). These results are in agreement with the theoretical predictions in Fig. 5B, suggesting that for the given iSTDP profile (Fig. 5A) midrange time shifts (20 < Δ*t*_stim_ < 80 ms) and low frequencies (*f*_stim_ < 20 Hz) are more likely to produce favourable post-stimulation outcomes, i.e., reduce synaptic wiring as well as neuronal synchrony. In addition, increasing the stimulation amplitude promotes the effectiveness of stimulation in reducing STN-GPe synchrony and synaptic wiring (Fig. 10B), due to an increase in the strength of neuronal spike entrainment (also see Fig. 6A and B).

**Figure 10:**
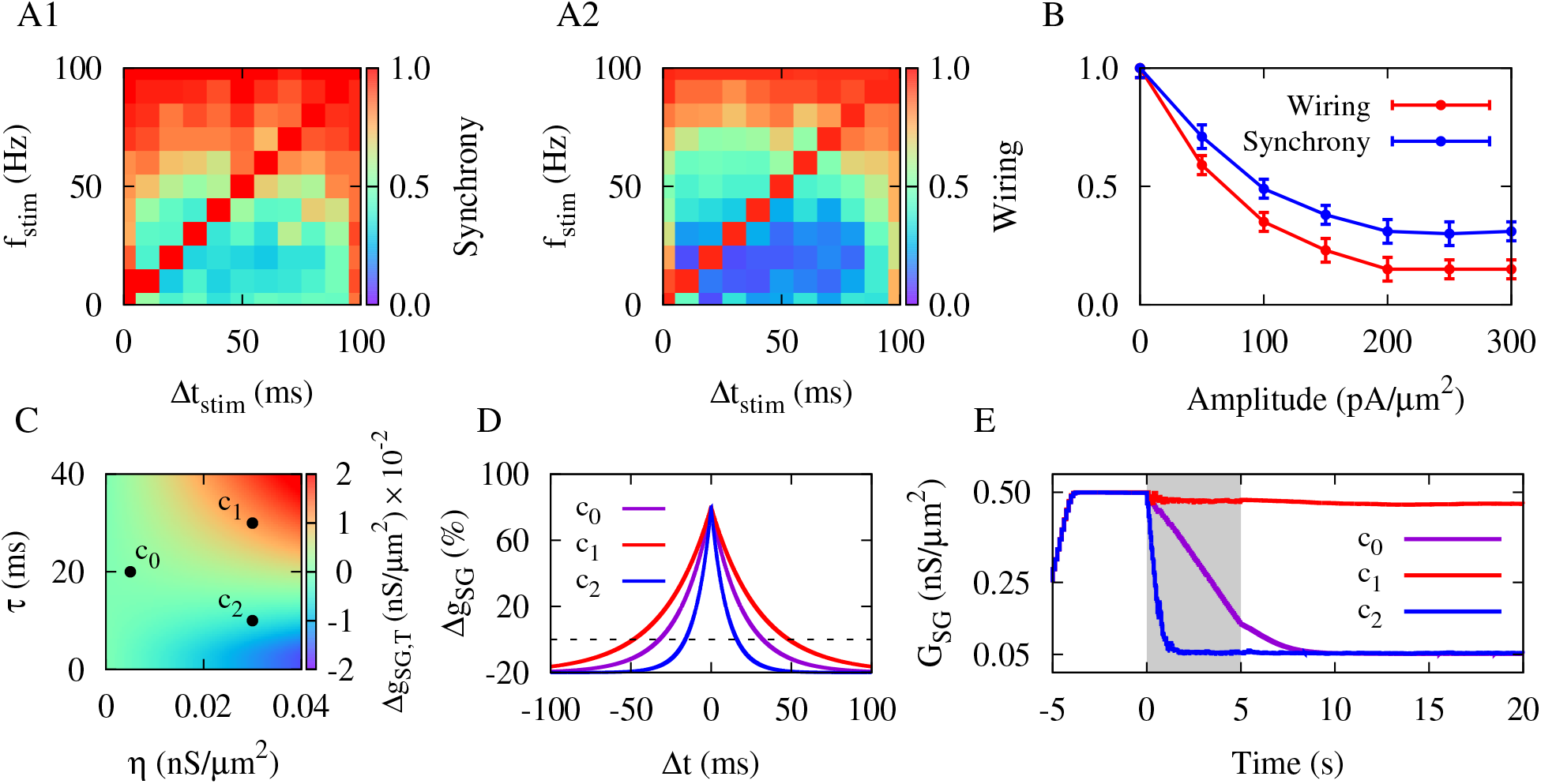
Re-tuning of stimulation parameters and plasticity profile. (**A1**,**A2**) The STN-GPe synchrony (A1) and synaptic wiring (A2) post-stimulation with different time shifts and frequencies. (**B**) The impact of stimulation amplitude on the STN-GPe synchrony and synaptic wiring. (**C**) Synaptic change based on Eq. (19) with red (blue) representing synaptic potentiation (depression) when the iSTDP learning rate and decay time constant were varied. Point c_0_ with (*η, τ*) = (0.005 nS/*µ*m^2^, 20 ms) was used as the default iSTDP profile, whereas point c_1_ with (0.03 nS/*µ*m^2^, 30 ms) and point c_2_ with (0.03 nS/*µ*m^2^, 10 ms) indicate tested iSTDP parameters. (**D**) The iSTDP profile described by Eq. (14) corresponding to three parameter sets marked in panel C. Dashed line indicates Δ*g*_SG_ = 0. (**E**) Time course of the mean GPe→ STN synaptic coupling for *c*_0_, *c*_1_ and *c*_2_ sets. The shaded area indicates the stimulation epoch (0 < *t* < 5 s). To be able to compare the mean couplings, the iSTDP parameters were identical for *t* < 0 (with parameters given by *c*_0_) and then stayed the same (*c*_0_) or were switched to new values (*c*_1_ and *c*_2_) at *t* = 0 for the rest of simulation.

Nonetheless, for a given set of stimulation time shift and frequency values, variations of the iSTDP profile may significantly influence the ability of the rhythmic stimulation paradigm to alter synaptic wiring and, consequently, affect neuronal synchrony. To explore the effects of the plasticity profile on model outcomes, we systematically varied the iSTDP learning rate (*η*) and decay time constant (*τ*) in Fig. 10C and calculated the resultant GPe→STN synaptic change based on Eq. (19). Fig. 10C shows that, depending on the iSTDP parameters, the synaptic weights may be weakened (blue) or strengthened (red).

We highlight three sets of parameters in Fig. 10C to compare model outcomes as a function of the plasticity parameters, i.e., point c_0_ with (*η, τ*) = (0.005 nS/*µ*m^2^, 20 ms) used as the default iSTDP profile throughout the manuscript, point c_1_ with (0.03 nS/*µ*m^2^, 30 ms) and point c_2_ with (0.03 nS/*µ*m^2^, 10 ms). The corresponding iSTDP profiles of the three sets of parameters are depicted in Fig. 10D. When the iSTDP profile is characterised by a greater potentiation regime compared to the default iSTDP profile (cf. Fig. 10D, red and violet), stimulation fails to successfully reduce the mean GPe→ STN synaptic coupling (cf. Fig. 10E, red and violet). However, for the iSTDP profile with a smaller potentiation regime compared to the default iSTDP profile (cf. Fig. 10D, blue and violet), stimulation reduces the mean synaptic coupling even faster than the default iSTDP profile (cf. Fig. 10E, blue and violet). Together, these results indicate that the shape of the plasticity learning window may crucially influence stimulation outcomes, demanding appropriate re-tuning of stimulation parameters based on model predictions.

## 4 Discussion

Non-invasive brain stimulation strategies that can provide long-lasting effects are a current clinical need for PD patients as they can induce sustained therapeutic after-effects. To that end,a stimulation protocol should be able to desynchronise aberrant synchronous neuronal activity and, furthermore, reduce pathologically strong synaptic connectivity to avoid relapse of symptoms. Importantly, even minor modifications of stimulation protocols may have unintended consequences on stimulation outcomes. We illustrate this by means of a simple STN-GPe loop with plastic synapses using computational modelling. More specifically, time-shifted dual STN-GPe stimulation led to a synaptic rewiring of the initially strong GPe→ STN connections followed by a suppression of the abnormally synchronised beta oscillations. Most importantly, this state remained stable and the pathological connectivity and excessive beta oscillations did not reappear even after external stimulation was discontinued. Interestingly, this principle works in a generic manner, no matter whether we use time-shifted continuous/intermittent stimulation, or monopolar/bipolar stimulation waveforms.

Stimulation-induced synaptic rewiring and control of synchronisation were previously investigated in several computational studies. For example, activation of multiple neuronal subpopulations with a time delay between different stimulation sites using a variety of temporal patterns can desynchronise oscillatory activity and may influence synaptic connectivity in adaptive neuronal networks [28,37,53,78]. Our study, on the other hand, emphasises the key role of plasticity in reshaping synaptic connections by stimulation and its implications for rehabilitation. In fact, the ability to modify synaptic connections becomes efficacious in disease [79–81], when connectivity and associated activity are compromised. Our rhythmic stimulation paradigm is based on two arguments. First, reshaping of the synaptic connections between two populations is a pairwise process entailing dual-site stimulation of both pre- and post-synaptic populations to engage synaptic plasticity. Second, an appropriate time shift between paired stimuli delivered to the two inter-connected populations is crucial to produce prolonged and robust changes in inter-population synaptic connections. Both arguments are in line with experimental observations in awake, behaving monkeys and computational models indicating that STDP could be induced via paired stimulation of two anatomically separate regions, depending on the time shift (Δ*t*_stim_) between stimuli [55, 56].

Stimulation protocols primarily aimed at desynchronising excessively synchronised activity may not necessarily decouple neurons during stimulation [27, 28]. Alternatively, long-lasting desynchronising effects can be realised by decoupling neurons both functionally and structurally, i.e., causing an unlearning of pathological interactions by simultaneously suppressing overly synchronised activity and reducing strong synaptic connectivity [30]. Plasticity-induced bistability of physiological and pathological model attractor states comprising the STN-GPe loop may facilitate this process [37, 42], allowing stimulation with appropriate parameters to shift activity and connectivity from a pathological model state (strong synchrony and strong connectivity) to a physiological model state (weak synchrony and weak connectivity) [27].

The frequency of stimulation is a key factor in achieving beneficial clinical outcomes. Traditionally, DBS is administered at high frequencies (> 100 Hz) for the treatment of movement disorders. For example, tremor is typically reduced by single-site stimulation at frequencies over 90 Hz [82–84], while motor symptoms are improved using stimulation frequencies below 60 Hz [85–87]. However, evidence suggests that relatively low-frequency stimulation may also be useful [88]. For instance, 60 Hz stimulation has been reported to improve axial parkinsonian symptoms such as gait freezing, swallowing, and speech difficulties [89–91]. Guided by theoretical predictions, our results show that in a dual-site stimulation model, an appropriate time shift between low-frequency stimuli may allow for pronounced, long-lasting changes in synaptic connectivity and oscillatory activity by harnessing synaptic plasticity, which may have potential therapeutic implications for this patient population.

The polarity of the stimulus waveform (cathodic vs. anodic) may determine which neural pathway is activated by stimulation, possibly engineering therapeutic benefits versus unwanted sideeffects [92, 93]. For example, as predicted by biophysical models and confirmed experimentally in PD patients [94], cathodic and anodic stimulation of the STN activate the same pathways in subcortical regions; however, cathodic stimulation is in general more efficient in activating the cortico-spinal and cortico-bulbar tracts as well as the cortico-subthalamic hyperdirect pathway. Yet, anodic stimulation is as efficient as cathodic in activating the STN-GPe pathway [94]. As shown in Fig. 9B and C, in our model, bipolar stimulation requires an extended stimulation ON epoch to be considered as efficient as monopolar stimulation.

We ignore transmission delays in the model. However, in principle, delays shape the emergent structure and dynamics in recurrent neuronal networks with plastic synapses by regulating synchronisation properties and synaptic connectivity [95–98]. In particular, inter-population delays in the STN-GPe loop are related to the latency of the cortex stimulation-induced responses in the basal ganglia downstream networks [99, 100], influencing healthy as well as parkinsonian oscillatory activity in computational models [101]. In our model, delays merely shift the effective temporal window of iSTDP required for the induction of synaptic potentiation/depression [96,97]. Therefore, the theoretical predictions of optimal stimulation time shift and frequency for effective synaptic rewiring must be re-tuned in the presence of transmission delays in the model [30].

Furthermore, we used a generic iSTDP profile to modify the synaptic strengths in the model, as shown in Fig. 5A. The temporally symmetric learning window of this iSTDP profile is a generic form of inhibitory synaptic plasticity observed in the hippocampal [39] and cortical [41] rodent slice preparations *in vitro*. However, the nature of long-term synaptic changes in the pre- and post-synaptically targeted STN and GPe populations has remained unknown, but could, in principle, be assessed experimentally. Nonetheless, the theoretical predictions in our model can be re-tuned according to any plasticity profile and the predicted results can be used as a guide for the rewiring of the STN-GPe network interactions by the time-shifted, dual-site stimulation, as demonstrated for the same iSTDP profile with different parameters (see Fig. 10).

Here, we focus on neural effects that persist even after the stimulation offset. These persistent effects are likely mediated by longer-lasting models of synaptic plasticity, such as STDP. Therefore, we neglected the potential role of short-term synaptic plasticity, which may modulate excitatory and inhibitory synapses differently, depending on the frequency and site of stimulation [47, 102–104]. For instance, it has been shown experimentally that electrical stimulation of thalamic structures with low frequencies (< 100 Hz) can induce glutamate-mediated short-term synaptic facilitation [104, 105], whereas GABA-mediated short-term synaptic depression is induced by STN stimulation [102, 103]. This site specificity of stimulation-induced short-term plasticity vanishes at higher frequencies (> 100 Hz), e.g., due to axonal and synaptic failure [106]. Importantly, considering only short-term or long-term synaptic plasticity may be insufficient to capture the full range of stimulation-induced effects in modelling studies, calling for integrated models for a more accurate description of the stimulation effects at the level of synapses as well as large neuronal networks [104, 107].

To demonstrate the concept of time-shifted, dual-site stimulation and to highlight the role of plasticity in stimulation-induced synaptic rewiring, we restricted our analysis to a minimal model of the STN-GPe loop interactions. In the model, the PD state was mimicked by tuning the strength of synaptic couplings and applied currents where cortical and striatal inputs to the STN-GPe loop were simplified as external currents and the contribution of other pathways was ignored. However, PD pathophysiology is complex and multi-systemic in which abnormal beta oscillations can emerge due to a variety of interactions such as changes in cortical inputs [108], inhibitory interactions between striatal medium spiny neurons [109], competition between feedback loops [110], imbalance between cortico-subthalamic and pallido-subthalamic inputs [35] and increased levels of striatal cholinergic drive [111], in addition to reciprocal STN-GPe interactions [9,10]. While we do not claim that our simple model can recreate all complex interactions that lead to pathological oscillatory beta activity in PD, it is a well-understood basic model capable of reproducing parkinsonian dynamics computationally [11, 12, 14, 42, 54], allowing us to examine the efficacy of the time-shifted, dual-site stimulation.

Although many patients with PD benefit greatly from DBS therapy, not all patients experience optimal outcomes [112]. The complex circuit mechanisms underlying PD contribute to the variability in DBS efficacy [113]. Typically, DBS targets specific nodes of this circuit, such as the STN or globus pallidus, in order to restore healthy function. However, tremor, bradykinesia, rigidity, and axial symptoms in patients with PD map to different brain regions or networks [113], entailing personalised stimulation protocols targeting symptom-specific networks [114]. Although our paradigm is restricted to a model of STN-GPe, it has broad clinical applicability.

Ultimately, our model aims to establish a conceptual foundation for the time-shifted stimulation of (multiple) anatomically separate neuronal populations which is applicable across different stimulation modalities targeting various brain networks non-invasively. For instance, applying dual-site 10 Hz transcranial alternating current stimulation (tACS) with different phase shifts to two different cortical regions in humans modulates stimulation-outlasting functional connectivity, as assessed by electroencephalography (EEG) [115]. Later, computational modelling revealed that STDP of inter-population synapses can account for these experimentally observed connectivity after-effects of dual-site tACS, depending on the phase shift [116]. Furthermore, coactivation-induced functional and structural cortical rewiring by sensory stimulation [117–119], for example, during perceptual performance, can be viewed as a form of plasticity-facilitated decoupling [120–122]. For example, synchronous co-activation of the digits in humans increases functional connectivity within primary somatosensory cortex due to the temporal coincidence between the two-digit inputs, but no significant change is induced following asynchronous co-activation, as evidenced by functional magnetic resonance imaging (fMRI) [118]. The mechanisms underlying these changes are not fully understood, yet, the strengthening/weakening of cortical synaptic connections through frequency-dependent synaptic plasticity may be a potential candidate linked to improvements in tactile discrimination performance [120–122], implying that stimulation-induced synaptic rewiring through plasticity might work in a rather generic way in the brain.

## 5 Conclusion

We demonstrate that long-lasting unlearning of pathological synaptic connectivity and desynchronisation effects can be achieved with appropriately time-shifted, dual-site stimulation. Such stimulation not only reduces synaptic weights during stimulation, but also sufficiently reshapes synaptic wiring patterns, so that after stimulation the targeted network comprising two interacting neuronal populations (i.e., STN and GPe) relaxes into a decoupled state (characterised by weak synaptic coupling and weak synchrony), without further stimulus administration. The results of our modelling study might have strong implications in a variety of bi-/multi-channel stimulation techniques that are designed to target plasticity mechanisms in the brain to restore healthy patterns of activity and connectivity.

## CRediT Author Statement

**Mojtaba Madadi Asl:** Conceptualisation, Methodology, Formal analysis, Visualisation, Investigation, Writing - original draft, Writing - review & editing, Project administration, Supervision. **Caroline A. Lea-Carnall:** Methodology, Formal analysis, Investigation, Writing - review & editing, Supervision.

## Declaration of Competing Interests

The authors declare that the research was conducted in the absence of any commercial or financial relationships that could be construed as a potential conflict of interest.

## Data Availability

All data used to produce the figures were generated via numerical simulations. The simulation code is publicly accessible at https://github.com/MMadadiAsl/STN-GPe-synaptic-rewiring.

## Notes

### Competing Interest Statement

The authors have declared no competing interest.

## References

[1] da Silva FHL, Blanes W, Kalitzin SN, Parra J, Suffczynski P, Velis DN. Dynamical diseases of brain systems: different routes to epileptic seizures. IEEE Transactions on Biomedical Engineering. 2003;50(5):540–548.

[2] Weisz N, Moratti S, Meinzer M, Dohrmann K, Elbert T. Tinnitus perception and distress is related to abnormal spontaneous brain activity as measured by magnetoencephalography. PLoS Medicine. 2005;2(6):e153.

[3] Uhlhaas PJ, Singer W. Neural synchrony in brain disorders: relevance for cognitive dysfunctions and pathophysiology. Neuron. 2006;52(1):155–168.

[4] Hammond C, Bergman H, Brown P. Pathological synchronization in Parkinson’s disease: networks, models and treatments. Trends in Neurosciences. 2007;30(7):357–364.

[5] Uhlhaas PJ, Singer W. Abnormal neural oscillations and synchrony in schizophrenia. Nature Reviews Neuroscience. 2010;11(2):100–113.

[6] Plenz D, Kital ST. A basal ganglia pacemaker formed by the subthalamic nucleus and external globus pallidus. Nature. 1999;400(6745):677–682.

[7] Bevan MD, Magill PJ, Terman D, Bolam JP, Wilson CJ. Move to the rhythm: oscillations in the subthalamic nucleus–external globus pallidus network. Trends in Neurosciences. 2002;25(10):525–531.

[8] Mallet N, Pogosyan A, Sharott A, Csicsvari J, Bolam JP, Brown P, et al. Disrupted dopamine transmission and the emergence of exaggerated beta oscillations in subthalamic nucleus and cerebral cortex. Journal of Neuroscience. 2008;28(18):4795–4806.

[9] Mallet N, Pogosyan A, Márton LF, Bolam JP, Brown P, Magill PJ. Parkinsonian beta oscillations in the external globus pallidus and their relationship with subthalamic nucleus activity. Journal of Neuroscience. 2008;28(52):14245–14258.

[10] Tachibana Y, Iwamuro H, Kita H, Takada M, Nambu A. Subthalamo-pallidal interactions underlying parkinsonian neuronal oscillations in the primate basal ganglia. European Journal of Neuroscience. 2011;34(9):1470–1484.

[11] Terman D, Rubin JE, Yew A, Wilson C. Activity patterns in a model for the subthalamopallidal network of the basal ganglia. Journal of Neuroscience. 2002;22(7):2963–2976.

[12] Holgado AJN, Terry JR, Bogacz R. Conditions for the generation of beta oscillations in the subthalamic nucleus–globus pallidus network. Journal of Neuroscience. 2010;30(37):12340–12352.

[13] Pavlides A, John Hogan S, Bogacz R. Improved conditions for the generation of beta oscillations in the subthalamic nucleus–globus pallidus network. European Journal of Neuroscience. 2012;36(2):2229–2239.

[14] Shouno O, Tachibana Y, Nambu A, Doya K. Computational model of recurrent subthalamo-pallidal circuit for generation of parkinsonian oscillations. Frontiers in Neuroanatomy. 2017;11:21.

[15] Kühn AA, Tsui A, Aziz T, Ray N, Brücke C, Kupsch A, et al. Pathological synchronisation in the subthalamic nucleus of patients with Parkinson’s disease relates to both bradykinesia and rigidity. Experimental Neurology. 2009;215(2):380–387.

[16] Neumann WJ, Staub-Bartelt F, Horn A, Schanda J, Schneider GH, Brown P, et al. Long term correlation of subthalamic beta band activity with motor impairment in patients with Parkinson’s disease. Clinical Neurophysiology. 2017;128(11):2286–2291.

[17] Kühn AA, Kupsch A, Schneider GH, Brown P. Reduction in subthalamic 8–35 Hz oscillatory activity correlates with clinical improvement in Parkinson’s disease. European Journal of Neuroscience. 2006;23(7):1956–1960.

[18] Weinberger M, Mahant N, Hutchison WD, Lozano AM, Moro E, Hodaie M, et al. Beta oscillatory activity in the subthalamic nucleus and its relation to dopaminergic response in Parkinson’s disease. Journal of Neurophysiology. 2006;96(6):3248–3256.

[19] Meissner W, Leblois A, Hansel D, Bioulac B, Gross CE, Benazzouz A, et al. Subthalamic high frequency stimulation resets subthalamic firing and reduces abnormal oscillations. Brain. 2005;128(10):2372–2382.

[20] Kühn AA, Kempf F, Brücke C, Doyle LG, Martinez-Torres I, Pogosyan A, et al. Highfrequency stimulation of the subthalamic nucleus suppresses oscillatory β activity in patients with Parkinson’s disease in parallel with improvement in motor performance. Journal of Neuroscience. 2008;28(24):6165–6173.

[21] Bronte-Stewart H, Barberini C, Koop MM, Hill BC, Henderson JM, Wingeier B. The STN beta-band profile in Parkinson’s disease is stationary and shows prolonged attenuation after deep brain stimulation. Experimental Neurology. 2009;215(1):20–28.

[22] Benabid AL, Chabardes S, Mitrofanis J, Pollak P. Deep brain stimulation of the subthalamic nucleus for the treatment of Parkinson’s disease. The Lancet Neurology. 2009;8(1):67–81.

[23] McIntyre CC, Savasta M, Kerkerian-Le Goff L, Vitek JL. Uncovering the mechanism (s) of action of deep brain stimulation: activation, inhibition, or both. Clinical Neurophysiology. 2004;115(6):1239–1248.

[24] Temperli P, Ghika J, Villemure JG, Burkhard P, Bogousslavsky J, Vingerhoets F. How do parkinsonian signs return after discontinuation of subthalamic DBS? Neurology. 2003;60(1):78–81.

[25] Baizabal-Carvallo JF, Jankovic J. Movement disorders induced by deep brain stimulation. Parkinsonism & Related Disorders. 2016;25:1–9.

[26] Tass PA. A model of desynchronizing deep brain stimulation with a demand-controlled coordinated reset of neural subpopulations. Biological Cybernetics. 2003;89(2):81–88.

[27] Tass PA, Majtanik M. Long-term anti-kindling effects of desynchronizing brain stimulation: a theoretical study. Biological Cybernetics. 2006;94(1):58–66.

[28] Kromer JA, Tass PA. Long-lasting desynchronization by decoupling stimulation. Physical Review Research. 2020;2(3):033101.

[29] Kromer JA, Tass PA. Synaptic reshaping of plastic neuronal networks by periodic multichannel stimulation with single-pulse and burst stimuli. PLoS Computational Biology. 2022;18(11):e1010568.

[30] Madadi Asl M, Valizadeh A, Tass PA. Decoupling of interacting neuronal populations by time-shifted stimulation through spike-timing-dependent plasticity. PLoS Computational Biology. 2023;19(2):e1010853.

[31] Moran RJ, Mallet N, Litvak V, Dolan RJ, Magill PJ, Friston KJ, et al. Alterations in brain connectivity underlying beta oscillations in Parkinsonism. PLoS Computational Biology. 2011;7(8):e1002124.

[32] Galvan A, Devergnas A, Wichmann T. Alterations in neuronal activity in basal ganglia-thalamocortical circuits in the parkinsonian state. Frontiers in Neuroanatomy. 2015;9:5.

[33] Madadi Asl M, Vahabie AH, Valizadeh A, Tass PA. Spike-timing-dependent plasticity mediated by dopamine and its role in Parkinson’s disease pathophysiology. Frontiers in Network Physiology. 2022;2(817524):1–18.

[34] Fan KY, Baufreton J, Surmeier DJ, Chan CS, Bevan MD. Proliferation of external globus pallidus-subthalamic nucleus synapses following degeneration of midbrain dopamine neurons. Journal of Neuroscience. 2012;32(40):13718–13728.

[35] Chu HY, Atherton JF, Wokosin D, Surmeier DJ, Bevan MD. Heterosynaptic regulation of external globus pallidus inputs to the subthalamic nucleus by the motor cortex. Neuron. 2015;85(2):364–376.

[36] Kumar A, Cardanobile S, Rotter S, Aertsen A. The role of inhibition in generating and controlling Parkinson’s disease oscillations in the basal ganglia. Frontiers in Systems Neuroscience. 2011;5:86.

[37] Lourens MA, Schwab BC, Nirody JA, Meijer HG, van Gils SA. Exploiting pallidal plasticity for stimulation in Parkinson’s disease. Journal of Neural Engineering. 2015;12(2):026005.

[38] Lindi SA, Mallet NP, Leblois A. Synaptic changes in pallidostriatal circuits observed in the parkinsonian model triggers abnormal beta synchrony with accurate spatio-temporal properties across the basal ganglia. Journal of Neuroscience. 2024;44(9).

[39] Woodin MA, Ganguly K, Poo Mm. Coincident pre-and postsynaptic activity modifies GABAergic synapses by postsynaptic changes in Cltransporter activity. Neuron. 2003;39(5):807–820.

[40] Vogels TP, Froemke RC, Doyon N, Gilson M, Haas JS, Liu R, et al. Inhibitory synaptic plasticity: spike timing-dependence and putative network function. Frontiers in Neural Circuits. 2013;7:119.

[41] D’amour JA, Froemke RC. Inhibitory and excitatory spike-timing-dependent plasticity in the auditory cortex. Neuron. 2015;86(2):514–528.

[42] Madadi Asl M, Asadi A, Enayati J, Valizadeh A. Inhibitory spike-timing-dependent plasticity can account for pathological strengthening of pallido-subthalamic synapses in Parkinson’s disease. Frontiers in Physiology. 2022;13(915626):1–13.

[43] Maistrenko YL, Lysyansky B, Hauptmann C, Burylko O, Tass PA. Multistability in the Kuramoto model with synaptic plasticity. Physical Review E. 2007;75(6):066207.

[44] Madadi Asl M, Valizadeh A, Tass PA. Delay-induced multistability and loop formation in neuronal networks with spike-timing-dependent plasticity. Scientific Reports. 2018;8(12068):1–15.

[45] Lea-Carnall CA, Tanner LI, Montemurro MA. Noise-modulated multistable synapses in a Wilson-Cowan-based model of plasticity. Frontiers in Computational Neuroscience. 2023;17:1017075.

[46] Madadi Asl M, Ramezani Akbarabadi S. Delay-dependent transitions of phase synchronization and coupling symmetry between neurons shaped by spike-timing-dependent plasticity. Cognitive Neurodynamics. 2023;17(2):523–536.

[47] Awad MZ, Vaden RJ, Irwin ZT, Gonzalez CL, Black S, Nakhmani A, et al. Subcortical short-term plasticity elicited by deep brain stimulation. Annals of Clinical and Translational Neurology. 2021;8(5):1010–1023.

[48] Maith O, Baladron J, Einhäuser W, Hamker FH. Exploration behavior after reversals is predicted by STN-GPe synaptic plasticity in a basal ganglia model. iScience. 2023;26(5).

[49] Tass PA, Qin L, Hauptmann C, Dovero S, Bezard E, Boraud T, et al. Coordinated reset has sustained aftereffects in Parkinsonian monkeys. Annals of Neurology. 2012;72(5):816–820.

[50] Adamchic I, Hauptmann C, Barnikol UB, Pawelczyk N, Popovych O, Barnikol TT, et al. Coordinated reset neuromodulation for Parkinson’s disease: proof-of-concept study. Movement Disorders. 2014;29(13):1679–1684.

[51] Wang J, Nebeck S, Muralidharan A, Johnson MD, Vitek JL, Baker KB. Coordinated reset deep brain stimulation of subthalamic nucleus produces long-lasting, dose-dependent - motor improvements in the 1-methyl-4-phenyl-1, 2, 3, 6-tetrahydropyridine non-human primate model of parkinsonism. Brain Stimulation. 2016;9(4):609–617.

[52] Bore JC, Campbell BA, Cho H, Pucci F, Gopalakrishnan R, Machado AG, et al. Long-lasting effects of subthalamic nucleus coordinated reset deep brain stimulation in the non-human primate model of parkinsonism: A case report. Brain Stimulation. 2022;15(3):598–600.

[53] Ebert M, Hauptmann C, Tass PA. Coordinated reset stimulation in a large-scale model of the STN-GPe circuit. Frontiers in Computational Neuroscience. 2014;8:154.

[54] McLoughlin C, Lowery M. Impact of network topology on neural synchrony in a model of the subthalamic nucleus-globus pallidus circuit. IEEE Transactions on Neural Systems and Rehabilitation Engineering. 2024;32:282–292.

[55] Seeman SC, Mogen BJ, Fetz EE, Perlmutter SI. Paired stimulation for spike-timing-dependent plasticity in primate sensorimotor cortex. Journal of Neuroscience. 2017;37(7):1935–1949.

[56] Shupe L, Fetz E. An integrate-and-fire spiking neural network model simulating artificially induced cortical plasticity. eNeuro. 2021;8(2).

[57] Oorschot DE. Total number of neurons in the neostriatal, pallidal, subthalamic, and substantia nigral nuclei of the rat basal ganglia: a stereological study using the cavalieri and optical disector methods. Journal of Comparative Neurology. 1996;366(4):580–599.

[58] Steiner LA, Tomás FJB, Planert H, Alle H, Vida I, Geiger JR. Connectivity and dynamics underlying synaptic control of the subthalamic nucleus. Journal of Neuroscience. 2019;39(13):2470–2481.

[59] Kita H, Kitai S. Intracellular study of rat globus pallidus neurons: membrane properties and responses to neostriatal, subthalamic and nigral stimulation. Brain Research. 1991;564(2):296–305.

[60] Kita H, Kita S. The morphology of globus pallidus projection neurons in the rat: an intracellular staining study. Brain Research. 1994;636(2):308–319.

[61] Sadek AR, Magill PJ, Bolam JP. A single-cell analysis of intrinsic connectivity in the rat globus pallidus. Journal of Neuroscience. 2007;27(24):6352–6362.

[62] Baufreton J, Kirkham E, Atherton JF, Menard A, Magill PJ, Bolam JP, et al. Sparse but selective and potent synaptic transmission from the globus pallidus to the subthalamic nucleus. Journal of Neurophysiology. 2009;102(1):532–545.

[63] Bahuguna J, Sahasranamam A, Kumar A. Uncoupling the roles of firing rates and spike bursts in shaping the STN-GPe beta band oscillations. PLoS Computational Biology. 2020;16(3):e1007748.

[64] Rubin JE, Terman D. High frequency stimulation of the subthalamic nucleus eliminates pathological thalamic rhythmicity in a computational model. Journal of Computational Neuroscience. 2004;16(3):211–235.

[65] Mazzoni A, Lindén H, Cuntz H, Lansner A, Panzeri S, Einevoll GT. Computing the local field potential (LFP) from integrate-and-fire network models. PLoS Computational Biology. 2015;11(12):e1004584.

[66] Vogels TP, Sprekeler H, Zenke F, Clopath C, Gerstner W. Inhibitory plasticity balances excitation and inhibition in sensory pathways and memory networks. Science. 2011;334(6062):1569–1573.

[67] Golomb D, Rinzel J. Clustering in globally coupled inhibitory neurons. Physica D: Nonlinear Phenomena. 1994;72(3):259–282.

[68] Lachaux JP, Rodriguez E, Martinerie J, Varela FJ. Measuring phase synchrony in brain signals. Human Brain Mapping. 1999;8(4):194–208.

[69] Lemos JC, Friend DM, Kaplan AR, Shin JH, Rubinstein M, Kravitz AV, et al. Enhanced GABA transmission drives bradykinesia following loss of dopamine D2 receptor signaling. Neuron. 2016;90(4):824–838.

[70] Chu HY, McIver EL, Kovaleski RF, Atherton JF, Bevan MD. Loss of hyperdirect pathway cortico-subthalamic inputs following degeneration of midbrain dopamine neurons. Neuron. 2017;95(6):1306–1318.

[71] Kita H, Kita T. Role of striatum in the pause and burst generation in the globus pallidus of 6-OHDA-treated rats. Frontiers in Systems Neuroscience. 2011;5:42.

[72] Gerstner W, Kempter R, van Hemmen JL, Wagner H. A neuronal learning rule for sub-millisecond temporal coding. Nature. 1996;383(6595):76.

[73] Markram H, Lübke J, Frotscher M, Sakmann B. Regulation of synaptic efficacy by coincidence of postsynaptic APs and EPSPs. Science. 1997;275(5297):213–215.

[74] Bi GQ, Poo MM. Synaptic modifications in cultured hippocampal neurons: dependence on spike timing, synaptic strength, and postsynaptic cell type. Journal of Neuroscience. 1998;18(24):10464–10472.

[75] Brocker DT, Swan BD, So RQ, Turner DA, Gross RE, Grill WM. Optimized temporal pattern of brain stimulation designed by computational evolution. Science Translational Medicine. 2017;9(371):eaah3532.

[76] Merrill DR, Bikson M, Jefferys JG. Electrical stimulation of excitable tissue: design of efficacious and safe protocols. Journal of Neuroscience Methods. 2005;141(2):171–198.

[77] Popovych OV, Tass PA. Adaptive delivery of continuous and delayed feedback deep brain stimulation-a computational study. Scientific Reports. 2019;9(1):10585.

[78] Schmalz J, Kumar G. Controlling synchronization of spiking neuronal networks by harnessing synaptic plasticity. Frontiers in Computational Neuroscience. 2019;13:61.

[79] Bunday KL, Perez MA. Motor recovery after spinal cord injury enhanced by strengthening corticospinal synaptic transmission. Current Biology. 2012;22(24):2355–2361.

[80] Van Hartevelt TJ, Cabral J, Deco G, Møller A, Green AL, Aziz TZ, et al. Neural plasticity in human brain connectivity: the effects of long term deep brain stimulation of the subthalamic nucleus in Parkinson’s disease. PLoS ONE. 2014;9(1):e86496.

[81] Rebelo D, Oliveira F, Abrunhosa A, Januário C, Lemos J, Castelo-Branco M. A link between synaptic plasticity and reorganization of brain activity in Parkinson’s disease. Proceedings of the National Academy of Sciences. 2021;118(3):e2013962118.

[82] Ushe M, Mink JW, Revilla FJ, Wernle A, Schneider Gibson P, McGee-Minnich L, et al. Effect of stimulation frequency on tremor suppression in essential tremor. Movement Disorders. 2004;19(10):1163–1168.

[83] Fogelson N, Kühn AA, Silberstein P, Limousin PD, Hariz M, Trottenberg T, et al. Frequency dependent effects of subthalamic nucleus stimulation in Parkinson’s disease. Neuroscience Letters. 2005;382(1-2):5–9.

[84] Kuncel AM, Cooper SE, Wolgamuth BR, Clyde MA, Snyder SA, Montgomery Jr EB, et al. Clinical response to varying the stimulus parameters in deep brain stimulation for essential tremor. Movement Disorders. 2006;21(11):1920–1928.

[85] Timmermann L, Wojtecki L, Gross J, Lehrke R, Voges J, Maarouf M, et al. Ten-Hertz stimulation of subthalamic nucleus deteriorates motor symptoms in Parkinson’s disease. Movement Disorders. 2004;19(11):1328–1333.

[86] Eusebio A, Chen CC, Lu CS, Lee ST, Tsai CH, Limousin P, et al. Effects of low-frequency stimulation of the subthalamic nucleus on movement in Parkinson’s disease. Experimental Neurology. 2008;209(1):125–130.

[87] Chen CC, Lin WY, Chan HL, Hsu YT, Tu PH, Lee ST, et al. Stimulation of the subthalamic region at 20 Hz slows the development of grip force in Parkinson’s disease. Experimental Neurology. 2011;231(1):91–96.

[88] Baizabal-Carvallo JF, Alonso-Juarez M. Low-frequency deep brain stimulation for movement disorders. Parkinsonism & Related Disorders. 2016;31:14–22.

[89] Moreau C, Defebvre L, Destée A, Bleuse S, Clement F, Blatt JL, et al. STN-DBS frequency effects on freezing of gait in advanced Parkinson disease. Neurology. 2008;71(2):80–84.

[90] Sidiropoulos C, Walsh R, Meaney C, Poon Y, Fallis M, Moro E. Low-frequency subthalamic nucleus deep brain stimulation for axial symptoms in advanced Parkinson’s disease. Journal of Neurology. 2013;260:2306–2311.

[91] Xie T, Vigil J, MacCracken E, Gasparaitis A, Young J, Kang W, et al. Low-frequency stimulation of STN-DBS reduces aspiration and freezing of gait in patients with PD. Neurology. 2015;84(4):415–420.

[92] Kirsch AD, Hassin-Baer S, Matthies C, Volkmann J, Steigerwald F. Anodic versus cathodic neurostimulation of the subthalamic nucleus: a randomized-controlled study of acute clinical effects. Parkinsonism & Related Disorders. 2018;55:61–67.

[93] Soh D, Ten Brinke TR, Lozano AM, Fasano A. Therapeutic window of deep brain stimulation using cathodic monopolar, bipolar, semi-bipolar, and anodic stimulation. Neuromodulation. 2019;22(4):451–455.

[94] Borgheai SB, Opri E, Isbaine F, Cole ER, Jafari Deligani R, Laxpati NG, et al. Neural pathway activation in the subthalamic region depends on stimulation polarity. medRxiv. 2024;p. 2024–05.

[95] Ernst U, Pawelzik K, Geisel T. Synchronization induced by temporal delays in pulse-coupled oscillators. Physical Review Letters. 1995;74(9):1570–1573.

[96] Lubenov EV, Siapas AG. Decoupling through synchrony in neuronal circuits with propagation delays. Neuron. 2008;58(1):118–131.

[97] Madadi Asl M, Valizadeh A, Tass PA. Dendritic and axonal propagation delays determine emergent structures of neuronal networks with plastic synapses. Scientific Reports. 2017;7(39682):1–12.

[98] Madadi Asl M, Valizadeh A, Tass PA. Dendritic and axonal propagation delays may shape neuronal networks with plastic synapses. Frontiers in Physiology. 2018;9(1849):1–8.

[99] Nambu A, Tokuno H, Hamada I, Kita H, Imanishi M, Akazawa T, et al. Excitatory cortical inputs to pallidal neurons via the subthalamic nucleus in the monkey. Journal of Neurophysiology. 2000;84(1):289–300.

[100] Kita H, Kita T. Cortical stimulation evokes abnormal responses in the dopamine-depleted rat basal ganglia. Journal of Neuroscience. 2011;31(28):10311–10322.

[101] Asadi A, Madadi Asl M, Valizadeh A, Perc M. Dynamics of parkinsonian oscillations mediated by transmission delays in a mean-field model of the basal ganglia. Frontiers in Cellular Neuroscience. 2024;18:1344149.

[102] Bugaysen J, Bar-Gad I, Korngreen A. The impact of stimulation induced short-term synaptic plasticity on firing patterns in the globus pallidus of the rat. Frontiers in Systems Neuroscience. 2011;5:16.

[103] Milosevic L, Kalia SK, Hodaie M, Lozano AM, Fasano A, Popovic MR, et al. Neuronal inhibition and synaptic plasticity of basal ganglia neurons in Parkinson’s disease. Brain. 2018;141(1):177–190.

[104] Milosevic L, Kalia SK, Hodaie M, Lozano AM, Popovic MR, Hutchison WD, et al. A theoretical framework for the site-specific and frequency-dependent neuronal effects of deep brain stimulation. Brain Stimulation. 2021;14(4):807–821.

[105] Milosevic L, Kalia SK, Hodaie M, Lozano AM, Popovic MR, Hutchison WD. Physiological mechanisms of thalamic ventral intermediate nucleus stimulation for tremor suppression. Brain. 2018;141(7):2142–2155.

[106] Rosenbaum R, Zimnik A, Zheng F, Turner RS, Alzheimer C, Doiron B, et al. Axonal and synaptic failure suppress the transfer of firing rate oscillations, synchrony and information during high frequency deep brain stimulation. Neurobiology of Disease. 2014;62:86–99.

[107] Farokhniaee A, McIntyre CC. Theoretical principles of deep brain stimulation induced synaptic suppression. Brain Stimulation. 2019;12(6):1402–1409.

[108] Mallet N, Ballion B, Le Moine C, Gonon F. Cortical inputs and GABA interneurons imbalance projection neurons in the striatum of parkinsonian rats. Journal of Neuroscience. 2006;26(14):3875–3884.

[109] McCarthy M, Moore-Kochlacs C, Gu X, Boyden E, Han X, Kopell N. Striatal origin of the pathologic beta oscillations in Parkinson’s disease. Proceedings of the National Academy of Sciences. 2011;108(28):11620–11625.

[110] Leblois A, Boraud T, Meissner W, Bergman H, Hansel D. Competition between feedback loops underlies normal and pathological dynamics in the basal ganglia. Journal of Neuroscience. 2006;26(13):3567–3583.

[111] Kondabolu K, Roberts EA, Bucklin M, McCarthy MM, Kopell N, Han X. Striatal cholinergic interneurons generate beta and gamma oscillations in the corticostriatal circuit and produce motor deficits. Proceedings of the National Academy of Sciences. 2016;113(22):E3159–E3168.

[112] Yin Z, Cao Y, Zheng S, Duan J, Zhou D, Xu R, et al. Persistent adverse effects following different targets and periods after bilateral deep brain stimulation in patients with Parkinson’s disease. Journal of the Neurological Sciences. 2018;393:116–127.

[113] McGregor MM, Nelson AB. Circuit mechanisms of Parkinson’s disease. Neuron. 2019;101(6):1042–1056.

[114] Rajamani N, Friedrich H, Butenko K, Dembek T, Lange F, Navrátil P, et al. Deep brain stimulation of symptom-specific networks in Parkinson’s disease. Nature Communications. 2024;15(1):4662.

[115] Schwab BC, Misselhorn J, Engel AK. Modulation of large-scale cortical coupling by transcranial alternating current stimulation. Brain Stimulation. 2019;12(5):1187–1196.

[116] Schwab BC, König P, Engel AK. Spike-timing-dependent plasticity can account for connectivity aftereffects of dual-site transcranial alternating current stimulation. NeuroImage. 2021;237:118179.

[117] Godde B, Stauffenberg B, Spengler F, Dinse HR. Tactile coactivation-induced changes in spatial discrimination performance. Journal of Neuroscience. 2000;20(4):1597–1604.

[118] Vidyasagar R, Folger SE, Parkes LM. Re-wiring the brain: increased functional connectivity within primary somatosensory cortex following synchronous co-activation. NeuroImage. 2014;92:19–26.

[119] Schmidt-Wilcke T, Wulms N, Heba S, Pleger B, Puts N, Glaubitz B, et al. Structural changes in brain morphology induced by brief periods of repetitive sensory stimulation. NeuroImage. 2018;165:148–157.

[120] Lea-Carnall CA, Trujillo-Barreto NJ, Montemurro MA, El-Deredy W, Parkes LM. Evidence for frequency-dependent cortical plasticity in the human brain. Proceedings of the National Academy of Sciences. 2017;114(33):8871–8876.

[121] Brickwedde M, Schmidt MD, Krüger MC, Dinse HR. 20 Hz steady-state response in somatosensory cortex during induction of tactile perceptual learning through LTP-like sensory stimulation. Frontiers in Human Neuroscience. 2020;14:257.

[122] Lea-Carnall CA, Williams SR, Sanaei-Nezhad F, Trujillo-Barreto NJ, Montemurro MA, El-Deredy W, et al. GABA modulates frequency-dependent plasticity in humans. iScience. 2020;23(11):101657.

